# TTC7A organizes glandular lumen formation in the intestine through a Class II phosphatidylinositol 3-kinase

**DOI:** 10.1101/2025.03.22.644724

**Authors:** Katlynn Bugda Gwilt, Dhanushan Wijayaratna, Geordie Emberling, Lily Gillette, Jeffrey La, Hengqi Betty Zheng, Danielle Wendel, Lina B Karam, Aleixo M Muise, Scott B Snapper, Wayne I Lencer, Krishnan Raghunathan, Jay R. Thiagarajah

## Abstract

The formation of a single central lumen is a critical step for glandular morphogenesis. Patients with loss of function variants in the chaperone protein TTC7A have multiple lumen formation in colonic crypt glands. We show that trafficking and localization of TTC7A to the plasma membrane is required for directionally specifying the apical membrane with TTC7A patient loss-of-function variants leading to mislocalized membrane and lumen formation. Our experiments show that TTC7A, in early stages of apical membrane development, functions as a molecular chaperone for the Class II phosphatidylinositol 3-kinase, PIK3C2A and is trafficked in Rab11a positive vesicles to generate phosphatidylinositol 3,4-bisphosphate (PI(3,4)P_2_). We show that the apical specification process is dependent on PIK3C2A dependent generation of PI(3,4)P_2_ in intestinal epithelia and that defective lumen formation can be rescued by exogenous PI(3,4)P_2_ or small molecules that modulate phosphoinositide homeostasis.

## Main

Glandular epithelial structures called crypts are a key morphological feature of the intestine. In the colon, most epithelial cells reside within these crypt glands and are critical for absorption, secretion, and intestinal barrier function. Glandular lumen formation requires initial polarization and specification of epithelial cells to form two distinct membrane domains: the lumen-facing apical membrane and the basolateral membrane, which interacts with neighboring cells and the extracellular matrix. Functional specialization of these membranes is enabled and maintained by their unique protein and lipid compositions^1^. The process of epithelial membrane organization and cell polarization have been extensively studied in a variety of model systems^2–7^. Studies using three-dimensional cyst models have provided evidence that polarity is determined early on during initial cell division, with formation of cell-cell contact sites enriched in phosphatidylinositol 4,5- bisphosphate (PI(4,5)P2)^2^ and in phosphatidylinositol 3,4-bisphosphate (PI(3,4)P2)^8^. Following enrichment of phosphoinositides (PIPs) and localization of specific polarity proteins^9–11^ at nascent opposing apical membranes, lumen formation occurs through directed vesicular trafficking pathways that deliver phosphoinostitide kinases, phosphatases, and other targeted proteins^2,7,8,12–14^.

TetraTriCopeptide repeat domain 7A (TTC7A) is a protein expressed in intestinal epithelial cells, with several lines of evidence suggesting a key role in cellular organization and survival. Loss of function variants in the gene encoding TTC7A result in a severe intestinal disease characterized by immune dysregulation, multiple intestinal atresia^15^, apoptotic enterocolitis^16^ and grossly abnormal epithelial architecture^17^. Previous studies have indicated loss of normal membrane polarity and increased cell death in patient derived intestinal enteroids^18^. In yeast, a homolog protein, TTC7B (ypp1) was identified as an important lipid modulator, chaperoning the phosphoinositide (PI) kinase (PIK) PI4KIIIα (stt4) to the plasma membrane in a complex with Efr3 and Fam126 (hycc1) to generate a plasma membrane pool of phosphatidylinositol 4-phosphate (PI4P)^19–21^. To date, most of the described functions of mammalian TTC7A are elucidated following studies of its yeast homolog TTC7B^22–25^. The role of human TTC7B as a PI4KIIIα scaffold was validated by crystallization of human TTC7B-PI4KIIIα-Fam126B^24^, and its function has been confirmed as a lipid kinase chaperone^22^. Corresponding interactions of TTC7A and PI4KIIIα have been observed patient derived immune cells and fibroblasts^26^ and in specific patient mutations in heterologous models^25^. suggest that TTC7A-dependent PI4KIIIα activity is important in regulating cellular apoptosis^18^. Patients with TTC7A loss of function, have profound alterations in colonic epithelial architecture with severe distortions of crypt and surface epithelial morphology.^27^ In these studies we sought to investigate how TTC7A regulates intestinal epithelial polarity and lumen formation and provide insight into the mechanisms underlying this fundamental developmental process.

## Results

### Mutations in TTC7A disrupt early lumen coalescence

Expression patterns of TTC7A and its homolog, TTC7B, differ across tissues and species. Given potentially distinct functional roles for TTC7A versus TTC7B and, since mutations in TTC7A not TTC7B, are known cause intestinal disease, we examined protein levels of TTC7A compared to TTC7B across intestinal tissues^28^. TTC7A is abundantly expressed throughout the intestine compared to TTC7B, which is only found at low levels in the rectum (Fig 1a). In colonic biopsies from healthy control pediatric patients there is robust expression of TTC7A protein in contrast to minimal and variable TTC7B expression (Fig 1b).

**Figure 1:**
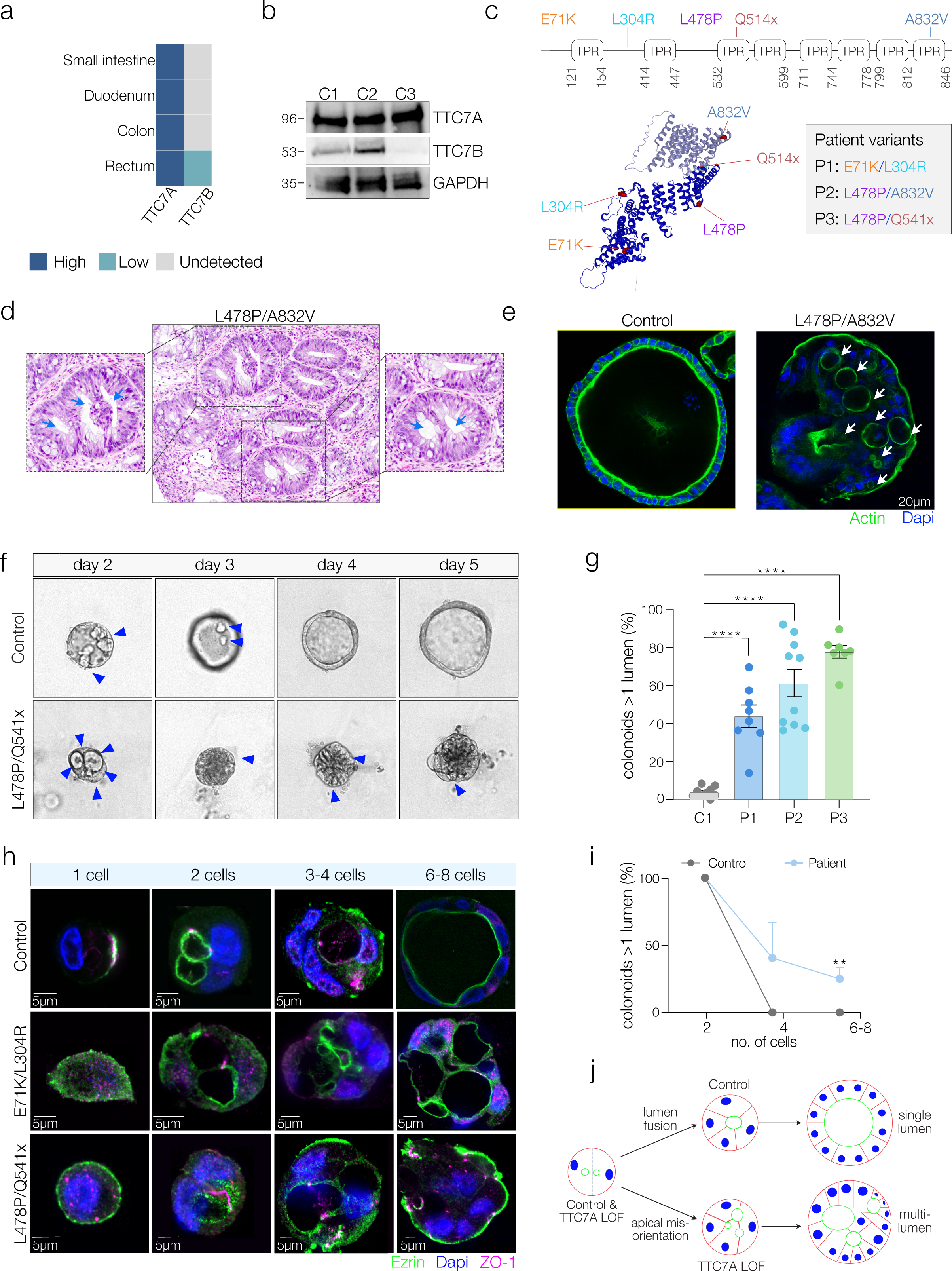
TTC7A mutations disrupt early lumen coalescence. **a.** TTC7A and TTC7B expression in small intestine, colon and rectum based on relative IHC expression from the human protein atlas. **b.** Western blot of TTC7A and TTC7B expression levels in three healthy colonic biopsies **c.** Visualization of TTC7A protein structure with compound heterozygous variants highlighted. **d.** Histopathology of patient tissue showing crypt glands with multiple secondary lumens in a patient with TTC7A mutations (L478P/A832V). Insets with blue arrows highlight secondary lumen structures in two distinct crypt glands. **e.** Representative confocal images of actin (green) and DAPI (blue) of healthy and TTC7A^LOF^ patient colonoids. **f.** Lumen formation assay by brightfield time-course imaging showing colon cyst development over 5 days, blue triangles mark secondary lumens. Representative of 3 separate experiments. **g.** Quantification of colonoids with more than one lumen in control vs. patient colonoids (n=3; data are shown as one-way ANOVA corrected for multiple comparisons; ****<0.0001). **h.** Representative confocal imaging of lumen development from single-cell through the 8-cell stage in control and patient-derived colonoids. Ezrin (green) and ZO-1 (red) at forming the apical lumen. **i.** Quantification of lumen coalescence by cell count in controls vs. patients (data representative of averages for n=3 patients and 3 controls). **j.** Working model illustrating secondary lumen coalescence in control colonoids and the loss of lumen coalescence in patient derived colonoids.

The crystal structure of TTC7A is experimentally unresolved, while the structure of human TTC7B has been partially crystalized in complexes with both Fam126A (PDB:5DSE^23^) and with Fam126B and PI4KIIIα (PDB:6BQ1^24^). Three sets of disease-causing TTC7A mutations from patients, were mapped onto the 3D Alphafold structure of TTC7A (Fig 1c). The compound heterozygous patients carried the following mutations: E71K/L304R, L478P/A832V, and L478P/Q541X. Despite the heterogeneous localization of mutations within the structure of TTC7A, all three patients exhibit a similar multi-lumen phenotype with disrupted epithelial architecture and crypt glands with secondary lumens apparent in colonic histology (Fig 1d and Supplementary 1a). To investigate the genesis of this phenotype, biopsy-derived colonoids were generated from all three patients. Cells with loss of TTC7A function are known to exhibit increased cell death in culture. To address this, culture conditions for long-term maintenance of TTC7A loss of function (TTC7A^LOF^) patient colonoids were optimized^29^, decreasing the percentage of death in all three patient lines (Supplementary 1a). Mature TTC7A^LOF^ colonoids consistently exhibited multiple secondary lumens as compared to mature colonoids from healthy controls reflective of the tissue phenotype. Additional lumens in TTC7A^LOF^ colonoids were lined with an actin rich brush border (Fig 1e) indicating extracellular lumen-facing apical membranes.

To track multi-lumen generation in colonoids, a time-course of early colonoid cyst formation was carried out using brightfield microscopy. Healthy control cells (Fig 1f, Supplementary 1b) consistently developed multiple lumens early in cyst development that resolved to a single central lumen by day 3 post-plating. In contrast TTC7A^LOF^ colonoids continued to exhibit secondary lumens at day 5 (Fig 1g) and beyond suggesting a failure of normal lumen coalescence seen in healthy controls. The secondary lumen structures present in mature TTC7A^LOF^ colonoids have expected polarized localization patterns of ezrin, ZO-1, and E-Cadherin at the apical membrane, tight junction, and basolateral domains (Supplementary 1c) respectively. Next, we conducted high-resolution fluorescence imaging of lumen coalescence from the single-cell stage to the 8+ cell stage (Fig 1h) by tracking early apical markers ezrin and ZO-1 along with actin. We observed the presence of secondary lumens in patients and controls at the two-cell stage (Fig 1h, 1i Supplementary 1d). These secondary lumens coalesce and are resolved during lumen expansion at the 3-4 cell stage in controls, but not in TTC7A patients (Fig 1i). These data suggest that multi-lumen formation in TTC7A^LOF^ colonoids occurs through the loss of normal coalescence early in colonoid development, likely due to mis-orientation and not loss of polarization of the lumen-facing apical membrane (Fig 1j).

### The localization of TTC7A changes during apical membrane specification

Given that TTC7A functions early in apical membrane organization, we followed TTC7A localization during cell polarization. LS174T:W4 (W4) cells are a colonic cell model^30^ that can be induced to form a single brush border-like structure rich in actin and villin (Fig 2a) following doxycycline administration allowing precise temporal control of colonic epithelial polarization (Supplementary 2a). Knockdown of TTC7A in W4 cells (TTC7A-KD), results in the generation of two or more actin and villin rich brush-border like structures (Fig 2a, 2b, 2c) mirroring the biopsy and colonoid lumen phenotype (Fig 2b). Given the previous described interactions of TTC7A and TTC7B with the phosphoinositide kinase, PI4KIIIα, we assessed TTC7A and PI4KIIIα localization in WT-W4 and TTC7A-KD W4 cells. In fully polarized WT-W4 cells and TTC7A-KD cells, PI4KIIIα is predominantly cytosolic with minimal localization at membrane actin caps (Supplementary 2b). To understand the dynamic changes in TTC7A and PI4KIIIΑ, we tracked their localization during the transition from unpolarized to fully polarized in control WT cells. The localization of TTC7A transitions from a diffuse cytosolic distribution in unpolarized cells, to cytosolic puncta in partially polarized cells, and ultimately to the nascent apical membrane in fully polarized cells (Fig 2d). In unpolarized and partially polarized states, TTC7A and PI4KIIIΑ are relatively poorly spatially colocalized following polarization (Fig 2d, 2e, Supplementary 2b). A similarly result in the localization of TTC7A and PI4KIIIΑ is observed in early formation of control colonoids. TTC7A and PI4KIIIΑ were localized to distinct separate puncta in the subapical region in early lumen formation (Fig 2f). This contrasts with mature colonoids where we observe significant colocalization between TTC7A and PI4KIIIΑ at the apical membrane (Fig 2g). Previous reports demonstrate a loss of TTC7A-PI4KIIIΑ binding in cell models overexpressing TTC7A mutants^25^ but endogenous TTC7A-PI4KIIIΑ complex formation has not been assessed patient derived cells. We observed that the different TTC7A^LOF^ patient-derived colonoids had variable expression of TTC7A and a small but consistent reduction in association between TTC7A and PI4KIIIΑ both during early and mature lumen formation (Fig 2h, Supplementary 2d). The divergent localization of TTC7A with PI4KIIIΑ in early apical membrane specification and the observation that TTC7A localizes to sub-apical puncta early during polarization, led us to investigate whether alternative functions of TTC7A outside of its association with PI4KIIIΑ could be be responsible for directing glandular lumen formation.

**Figure 2:**
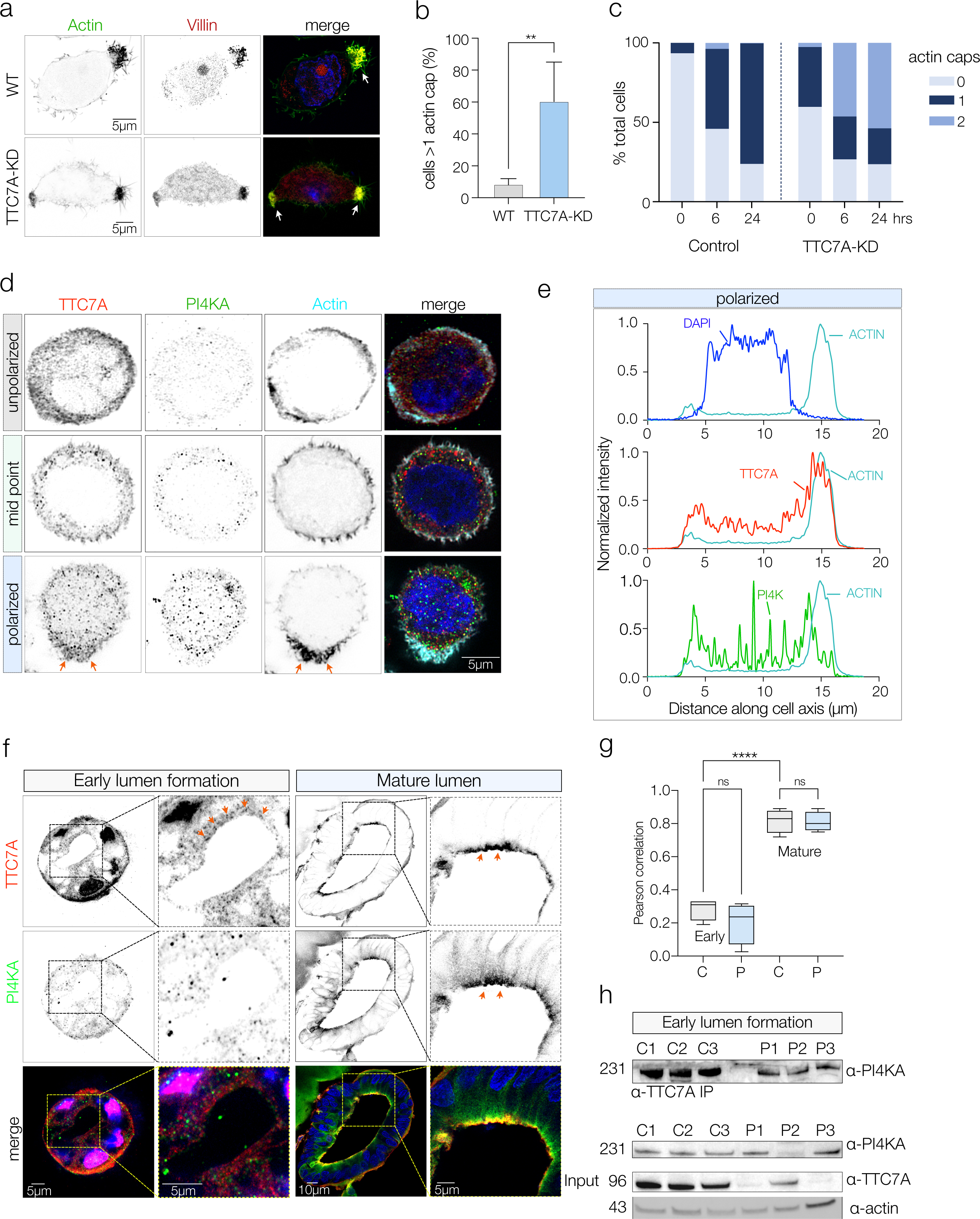
TTC7A localization changes during apical membrane specification. **a.** Representative confocal images with inverted micrographs showing immunofluorescent staining of actin caps (arrowheads) in LS174T:W4 control (WT) TTC7A-knockdown (KD) cells; actin (green) and villin (red), indicated by arrowheads. **b.** Quantification of secondary actin cap formation in TTC7A-KD cells. Mean of 3 experiments with 5 images per condition. (*p=<0.002). **c.** Quantification of actin cap formation across polarization in W4 cells; data represent two technical replicates, (n=5 images/time point). **d.** Representative confocal imaging with inverted micrographs showing endogenous localization of TTC7A (red) and PI4KIIIΑ (green) during W4 cell polarization; actin caps marked by red arrows. **e.** Representative plot profiles showing normalized intensity vs. cell length for the localization of Actin (teal) with DAPI (blue), TTC7A (red), and PI4KIIIΑ (green) at the endpoint of polarization. **f.** Representative confocal imaging of control colonoids showing endogenous TC7A (red) and PI4KIIIΑ (green) subcellular localization in early lumens and in mature lumens. Dashed boxes indicate the zoomed insets of the lumen. Insets show diverging TTC7A and PI4KIIIΑ in early lumen formation, and colocalization of TTC7A-PI4KIIIΑ in colonoids with a mature lumen by orange arrows. Grey-scale images are inverted black and white confocal images. **g.** Pearson’s correlation of TTC7A-PI4KIIIΑ colocalization in control vs. patient-derived colonoids. Data is representative of 4-6 experiments, One-way ANOVA, Tukey’s posthoc testing. ****<0.0001. **h.** Representative western blot on CO-IP TTC7A-pulldown probing for PI4KIIIΑ (top) in early lumen formation in three control and three patient colonoid lines. Data is representative of 6 experimental replicates. Western blot on whole cell lysates in three control and three patient colonoid lines probing for PI4KIIIΑ, TTC7A and actin. Data is representative of >6 experimental replicates.

### TTC7A^LOF^ is associated with decreased PI(3,4)P_2_ early during polarization

To investigate the function of TTC7A as a chaperone for lipid kinases, we first assessed the generation of the PI4KIIIΑ product PI(4)P following TTC7A knockdown in W4 cells or in TTC7A^LOF^ colonoids. Phosphoinositide (PIP) generation in cells is highly dynamic, with the action of both kinases and phosphatases leading to the same PIP species (Fig 3a). Using the PI(4)P lipid probe EGFP-P4M-SiDM^31^, we found PI4P levels were not significantly changed in TTC7A-KD W4 cells compared to WT-controls (Fig 3b) and were localized throughout the cell. This is not entirely unexpected given the plasticity of PI(4)P generation via multiple PI4-kinases (Fig 3a). Next, we assessed the key apical membrane phosphoinositides, PI(3,4)P_2_ and PI(4,5)P_2_. using GFP-C1- PHCdelta-PH^32^ which selectively binds PI(4,5)P_2_, or NES-EGFP-cPHx3^33^ that selectively binds PI(3,4)P_2_ . In TTC7A-KD cells, levels of PI(4,5)P_2_ were not significantly altered compared to control (Fig 3c). There was, however, a significantly reduced levels of PI(3,4)P_2_ in TTC7A-KD W4 cells (Fig 3d). We then validated our findings in control and patient-derived colonoids (Figure 3e-j). There were no significant changes in PI(4)P levels in TTC7A^LOF^ colonoids (Fig 3e, Supplementary 3a, 3b), but PI(3,4)P_2_ expression was significantly decreased in both mature (Fig 3e, 3f) and early forming colonoids (Fig 3g). There was no change in abundance of PI(4,5)P_2_ or changes in colocalization with TTC7A in control colonoids (Supplementary 3c, 3d), antithetical to any potential role for TTC7A to regulate PI(4,5)P_2_ during the early stages of lumen formation (Supplementary 3d). Conversely, PI(3,4)P_2_ is localized to juxta-membrane puncta during early lumen formation (Fig 3g, 3h), and is associated with TTC7A in control colonoids (Fig 3i, 3j) and is significantly reduced in TTC7A^LOF^ colonoids (Fig 3i, 3j). These results indicate that TTC7A has a potential function in regulating PI(3,4)P_2_ levels early in lumen formation.

**Figure 3:**
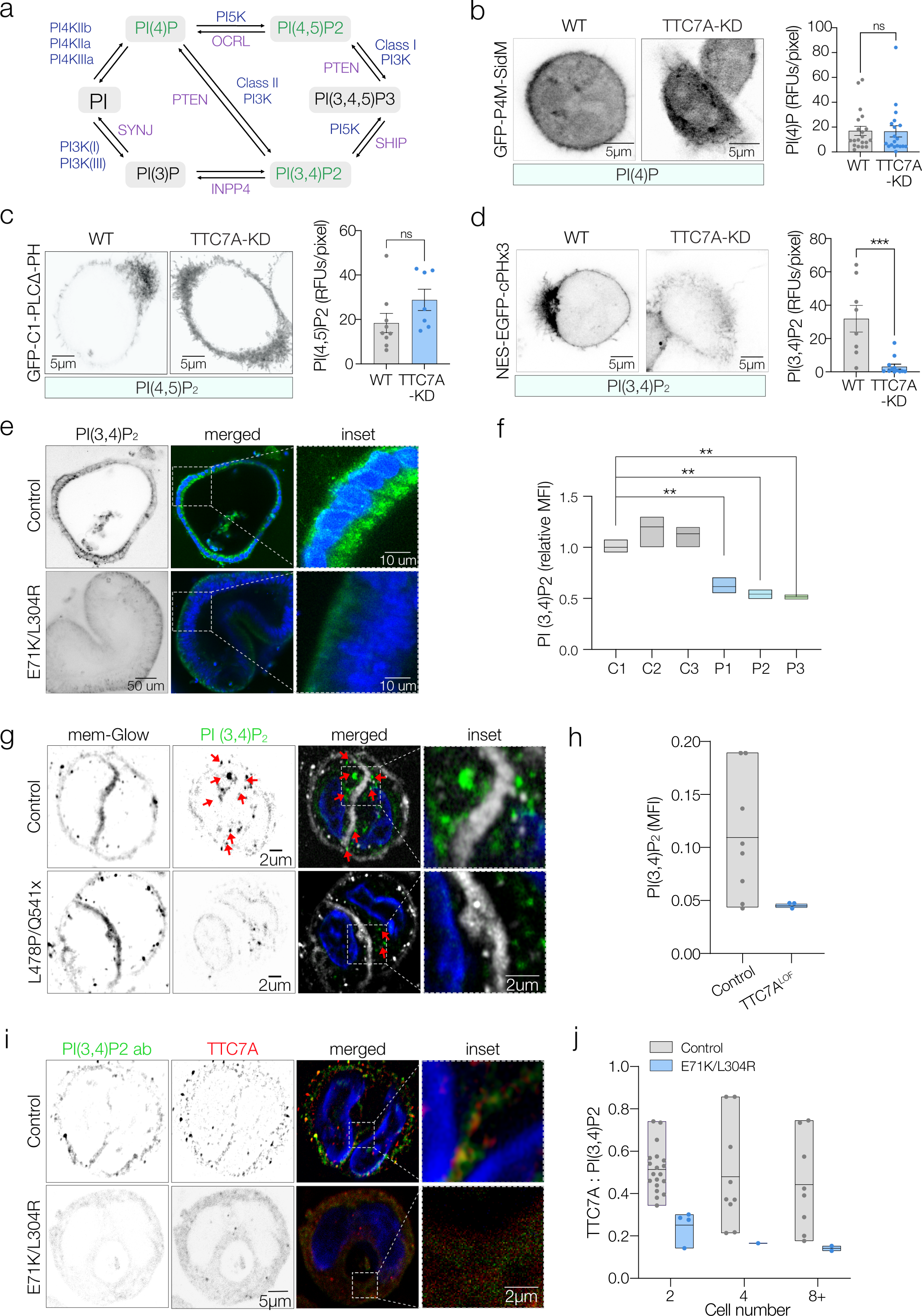
TTC7A^LOF^ is associated with decreased PI(3,4)P2 during polarization. **a**. Conversion between phosphoinositide species derived from PI occur by kinase action (blue) and phosphatase actions (purple). Two classes of PI4-kinase generate PI(4)P from PI. **b-d**. Representative confocal images with inverted micrographs to show the relative quantification of PI(4)P, PI(4,5)P2 and PI(3,4)P2 intensities in WT vs. TTC7A-KD cells expressing PI-GFP probes (GFP-P4M-SIDM, GFP-C1-PLCdelta-PH, and NES-EGFP-cPHx3 respectively). Data presented as RFUs per pixel; n=12+ replicates, unpaired two-tailed t-test, *<0.05, **<0.01, ***<0.001, ****<0.0001. **e.** Representative live confocal imaging on control and TTC7A^LOF^ colonoids showing PI(3,4)P2 probe subcellular localization (green) and nuclei (blue). **f.** Quantification of live confocal imaging for PI(3,4)P2 probe intensity across colonoid lines (n=3 controls, n=3 patients). Data presented as relative MFI. One-way ANOVA with Dunnett’s test *<0.05; **<0.01, ***<0.001, ****<0.0001. **g.** Representative confocal imaging with inverted micrographs of control and TTC7A^LOF^ colonoids at the two-cell state highlighting the localization of PI(34)P2 during de novo lumen formation. Membranes are marked by mem-glow (white) and PI(3,4)P2 (green). Dashed boxes are representative of the zoomed in inset at the cell-cell contact. Red arrows mark PI(3,4)P2 subcellular localization at the cell-cell contact. **h.** Quantification of PI(3,4)P2 intensity in control and TTC7A^LOF^ colonoids during the early stages of lumen formation. Data are represented as pooled populations of three controls and three TTC7A^LOF^ colonoids. **i.** Representative confocal imaging on endogenous TTC7A (red) and PI(3,4)P2 (green) in control and patient derived colonoids. Dashed box indicates zoomed inset with TTC7A and PI(3,4)P2 colocalization highlighted in yellow. **j.** Pearsons correlation of TTC7A: PI(3,4)P2 in control and TTC7A^LOF^colonoids in the 2-, 4-, and 8-cell stages of colonoid development.

### TTC7A associates with PIK3C2A in early lumen formation

Given our findings that loss of TTCA function is associated with altered levels of PI(3,4)P_2_, we reasoned that TTC7A could act as a chaperone for a corresponding lipid kinase to facilitate delivery through the cell. PI(3,4)P_2_ is a quantitatively minor phosphoinositide product in the cell generated through two processes: dephosphorylation of PI(3,4,5)P_3_ by SHIP1/2 or phosphorylation of PI(4)P by Class II PI-3 kinases notably, PIK3C2A (Fig 3a).^1^ In early-stage colonoids, we observed a population of TTC7A-PIK3C2A co-stained sub-membrane puncta resembling the pattern seen with PI(3,4)P_2_ (Fig 4a, 4b). Co-immunoprecipitation confirmed the interaction between TTC7A and PIK3C2A in control colonoids (Fig 4c). However, in both forming and mature TTC7A patient colonoids, there was a large and significant reduction in both colocalization and co-immunoprecipitation of TTC7A and PIK3C2A (Fig 4a, 4b, 4c, Supplementary 4a, 4b, 4c). Taken together with the previous observed loss of PI(3,4)P_2_ in TTC7A patients, these data suggest that in addition to its known role with PI4KIIIΑ^16^, TTC7A also acts as a chaperone for PIK3C2A to direct and localize generation of PI(3,4)P_2_ during apical membrane specification and lumenogenesis.

**Figure 4:**
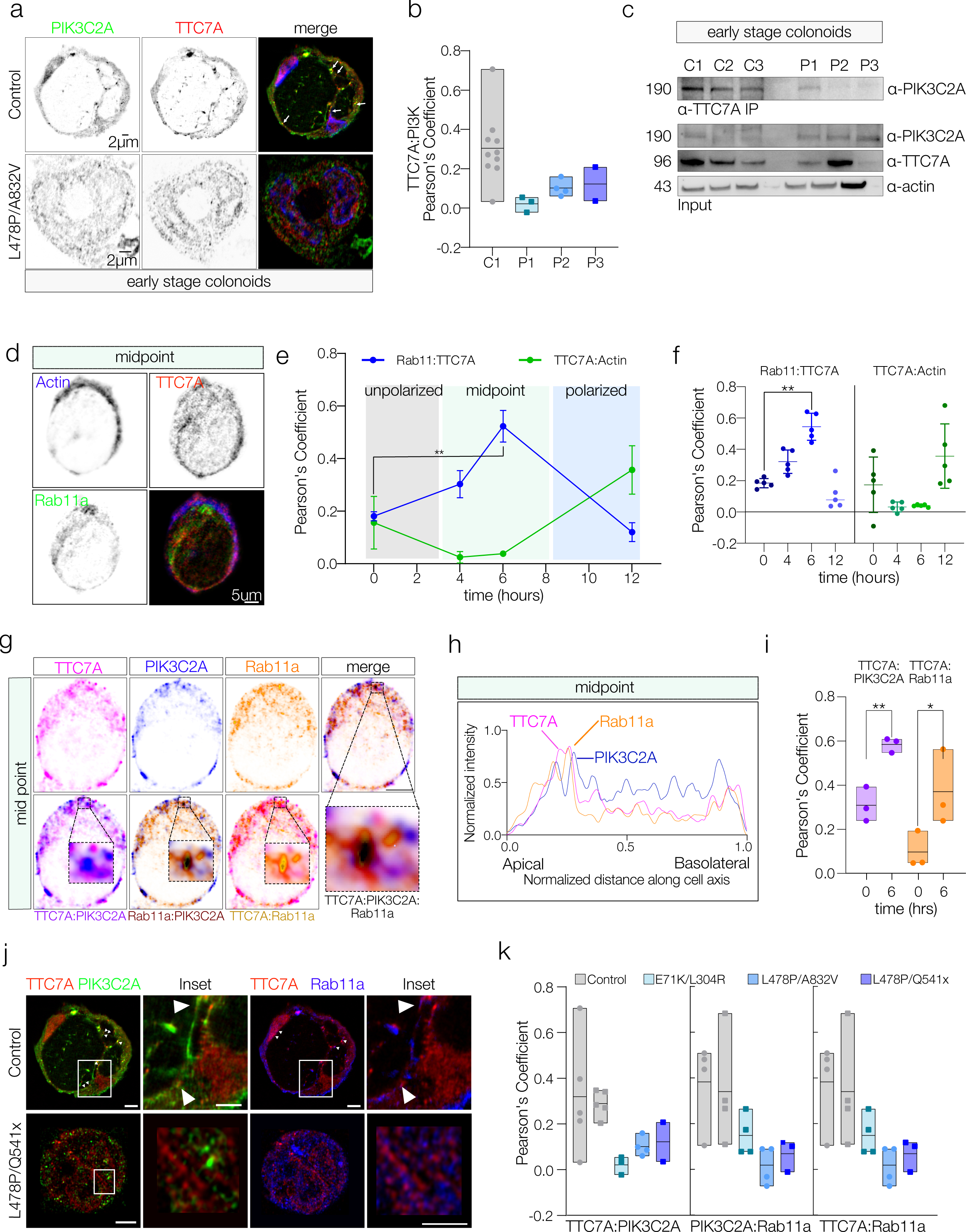
TTC7A associates with PIK3C2A and Rab11a in early lumen formation. **a.** Representative confocal imaging of PIK3C2A (green) and TTC7A (red) in early-stage control vs. TTC7A^LOF^ colonoids. Colocalization marked by white arrow heads. Scale bars = 2uM. **b.** Pearson’s correlation for TTC7A and PIK3C2A localization in early-stage control (C) and TTC7ALOF colonoids (P1, P2, P3). (n=5 images per sample, n=3 patients, n=3 controls). **c.** Representative western blot on CO-IP TTC7A-pulldown probing for PIK3C2A (top) in early lumen formation in three control and three patient colonoid lines. Data is representative of 6 experimental replicates. Western blot on whole cell lysates in three control and three patient colonoid lines probing for PIK3C2A, TTC7A and actin. Data is representative of >6 experimental replicates. **d.** Micrographs showing endogenous subcellular localization of Actin (blue) TTC7A (red) and Rab11a (green) at the midpoint of W4 cell polarization. Scale bars = 5 uM. **e.** Line plot for pearson’s Coefficient on WT-W4 for Rab11a:TTC7A (blue line) and TTC7A:Actin (green line) demonstrating average protein interactions across time, unpolarized (0-3hrs), midpoint (3-8hrs) and polarized (8-12hrs) cell states. **= p<0.001. One-way ANOVA with Dunnett’s multiple comparisons test. **f.** Dot plot for Pearson’s Coefficient for Rab11a:TTC7A (blue points) and TTC7A:Actin (green points) from 0-12hrs post polarization. Data representative of 5 experimental replicates, with 5 images per timepoint. **= p<0.001. One-way ANOVA with Dunnett’s multiple comparisons test. **g.** Inverted immunostaining at W4 polarization midpoint for TTC7A (pink), PIK3C2A (blue), and Rab11a (orange), and merged TTC7A:PIK3C2A:Rab11a showing colocalization (black), TTC7A:PIK3C2A colocalization (purple), TTC7A:Rab11a colocalization (yellow) and PIK3C2A:Rab11a colocalization (brown). Scale bars = 10um. **h.** Representative plot profile of midpoint from g, showing subapical colocalization of TTC7A (pink), Rab11a (orange) and PIK3C2A (blue). **i.** Dot plot for Pearson’s coefficient for TTC7A:PIK3C2A (purple) and TTC7A:Rab11a (orange) at 0 and 6 hours. **=p< *=p<0.05 . Unpaired, two tailed t-test. **j.** TTC7A (red) colocalization with PIK3C2A (green) and Rab11a (blue) in control vs. L478P/Q541x colonoids with PIK3C2A:TTC7A (green:red) and TTC7A:Rab11a (red:green). Arrows indicate areas of distinct colocalization. **k.** Pearson’s correlation of TTC7A:PIK3C2A, TTC7A:Rab11a and PIK3C2A:Rab11a in forming colonoids at the two-cell state (n=5 images per sample, n=3 patients, n=3 controls).

### TTC7A and PIK3C2A colocalize in Rab11a^+^ vesicles during apical membrane specification

Our data showing sub-apical membrane punctate localization of TTC7A during early lumen formation, suggested potential occupancy of a vesicular compartment. Previous studies have implicated Rab11a^+^ vesicles as key site containing class II PI3Ks involved in the generation of PI(3,4)P2 and the specification of the apical membrane^8^. To understand temporally the relationship of TTC7A with Rab11a, we quantified their co-localization in W4-cells during polarization. Here we found that TTC7A and Rab11a were highly colocalized at around 6 hours post induction which represents the midpoint of epithelial polarization in our model (Fig 4d, 4e, 4f Supp 4d) and not during the unpolarized or completely polarized state. Next, we quantified the analogous localization of PIK3C2A in the context of TTC7A and Rab11a in W4 cells over time (Fig 4g, Supplementary 4e). At all timepoints, TTC7A-PIK3C2A were more colocalized relative to TTC7A:Rab11a (Supplementary 4f, 4g). At the midpoint of epithelial polarization, TTC7A, PIK3C2A and Rab11a are primarily localized in a region proximal to the developing apical membrane (Fig 4g, 4h), with association of TTC7A with both PIK3C2A and Rab11a transiently increasing during formation of the apical membrane (Fig 4i). We validated these results by quantifying colocalization of TTC7A, PIK3C2A and Rab11a during early lumen formation in control and TTC7A^LOF^ colonoids. In control colonoids, we observed significantly higher colocalization of TTC7A, PIK3C2A and Rab11a in sub-membrane regions of cells (Fig 4j, 4k) compared to TTC7A^LOF^ colonoids. Taken together, our results suggests that chaperoning role of TTC7A during apical membrane specification likely proceeds through Rab11^+^ vesicles.

### Localization of TTC7A is critical for correct apical membrane specification

Our data in early forming colonoids and during polarization in W4 cells provide evidence for a model of lumen formation where TTC7A acts as a chaperone for PIK3C2A for directionally localizing PI(3,4)P_2_ to ensure apical membrane specification. To mechanistically test this model, we employed a chemo-genetic approach to control TTC7A localization using chemically induced dimerization between a TTC7A-FK506 binding protein (TTC7A-FKBP12) and FKBP-rapamycin binding protein fused to a Lyn plasma membrane localization signal (FRB-Lyn) (Fig 5a). Using this system, TTC7A can be reversibly recruited to the plasma membrane in W4 cells upon administration of rapamycin^34^ (Fig 5a). In combination with the temporal control of polarization state provided by the W4 cell system, this allowed us to study the importance of spatial localization of TTC7A in lumen formation (Fig 5b). Upon addition of doxycycline but in the absence of rapamycin, cells polarize, as previously characterized, with the formation of an actin-rich membrane in most cells over 6hrs (Fig 5c). Given that rapamycin is an established modulator of plasma membrane F-actin reorganization^35^, we tested whether rapamycin itself could stimulate actin cap formation in TTC7A-FKBP Lyn-FRB W4 cells. We observed that rapamycin treatment in W4 cells for 6 hours (Supplementary 5b) or TTC7A-FKBP Lyn-FRB overexpression by itself did not induce or alter actin cap formation (Fig 5c, Supplementary 5a), or the percentage of cells with multiple actin caps (Fig 5d, Supplementary 5c). To direct TTC7A, to the plasma membrane, TTC7A-FKBP/ Lyn-FRB cells were stimulated with doxycycline and then treated with rapamycin at 3hrs, representing the early midpoint, or at 5.5hrs when most cells had started forming actin caps (Fig 5e 5f, 5g, 5h). Treatment with rapamycin at 3 hours caused aberrant and increased actin localization throughout the plasma membrane (Fig 5e, 5f). Addition of rapamycin at the near polarized state (5.5hrs) in contrast had minimal effect on actin distribution in the membrane (Fig 5g, Supplementary 5d), or the presence of secondary actin caps (Fig 5h). In cells treated with rapamycin, TTC7a was colocalized with actin confirming induced localization at the plasma membrane (Fig 5i, 5j). Our findings indicate that both spatial and temporal control of TTC7A is required for apical membrane specification during epithelia polarization.

**Figure 5.**
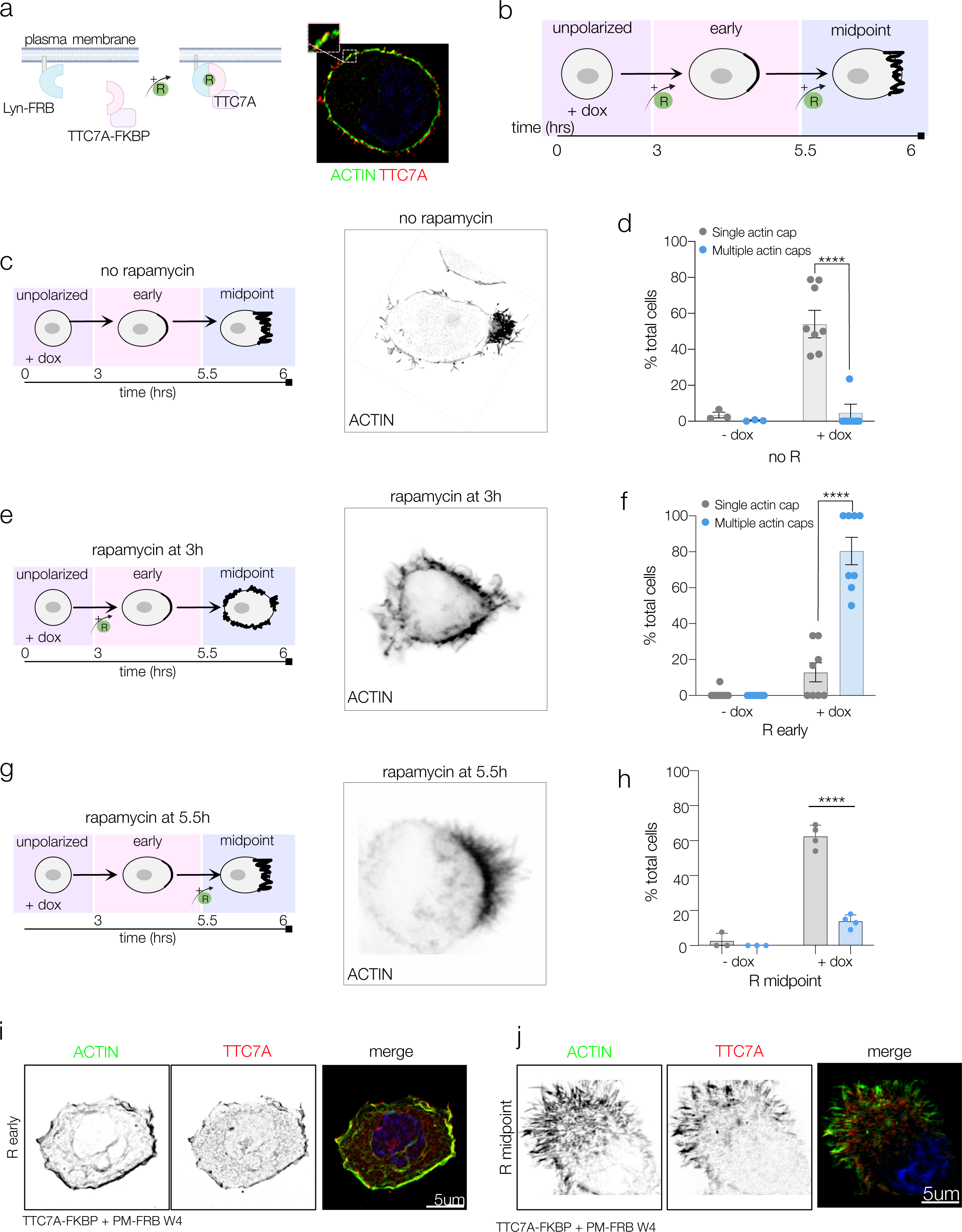
Apical membrane specification is dependent on TTC7A localization. **a.** TTC7A-FKBP and Lyn-FRB dimerization in W4 cells (TTC7A-FKBP PM-FRB) induced by rapamycin, with overexpression of TTC7A-FKBP (red) and endogenous expression of Actin (Green) shown in a non-polarized cell stimulated with rapamycin. Inset marks TTC7A:Actin colocalization (yellow) on the plasma membrane.. **b.** Schematic of the time course for doxycycline-induced polarization and rapamycin-induced dimerization in W4 cells treatment at 3 hours in the early-midpoint and 5.5 hours at the midpoint. **c.** Inverted black and white confocal image of actin cap formation of TTC7A-FKBP + PM-FRB W4 cells without rapamycin, visualized by actin staining (black and white). **d.** Quantification of baseline secondary actin cap formation in polarized TTC7A-FKBP + PM-FRB W4 cells. **e.** TTC7A-FKBP and Lyn-FRB induced at the early-midpoint of polarization, with actin staining (black and white). **f.** Quantification of multiple actin cap formation induced by rapamycin-mediated dimerization at the early-midpoint of polarization. **g.** TTC7A-FKBP and Lyn-FRB dimerization induced at the midpoint (5.5hrs post dox) with actin staining in black and white). **h.** Quantification of multiple actin cap formation with rapamycin-mediated dimerization at the midpoint of polarization. **i,j.** Immunostaining of actin (green) and TTC7A (red) in TTC7A-FKBP + Lyn-FRB W4 cells with dimerization induced either at the early midpoint or midpoint of polarization.

### Alteration of PIK3C2A and PI(3,4)P_2_ levels can recapitulate or rescue abnormal lumen formation in TTC7A disease

Given the evidence that TTC7A-dependent association and spatial localization of PIK3C2A is critical for directing apical membrane formation and lumen generation, we sought to investigate the effect of direct modulation of PIK3C2A function. Using siRNA constructs to knock down both PI4KIIIΑ and PIK3C2A in W4 cells (Supplementary 6a) we assessed actin cap formation. Loss of PIK3C2A led to a significant increase in secondary actin cap formation but not cell death (Fig 6a-c). In contrast loss of PI4KIIIΑ did not substantially increase secondary actin cap formation versus control cells (Supplementary 6b), but was accompanied by a significant increase in cell death (Supplementary 6e). To assess PIK3C2A function in colonoids, we treated healthy control colonoids with the specific PIK3C2A small molecule inhibitor PITCOIN3 ^36^. Loss of PIK3C2A enzymatic function induced by PITCOIN3 led to a substantial increase in multiple lumens in control colonoids, recapitulating the phenotype seen in TTC7A^LOF^ patients. (Fig 6d, 6e). In contrast inhibition of PI4KIIIΑ with the small molecule inhibitor GSKA1 did not significantly increase multi-lumen formation but, similar to loss of PI4KIIIΑ, did induce a significant increase in cell death (Supplementary 6c-e).

**Figure 6:**
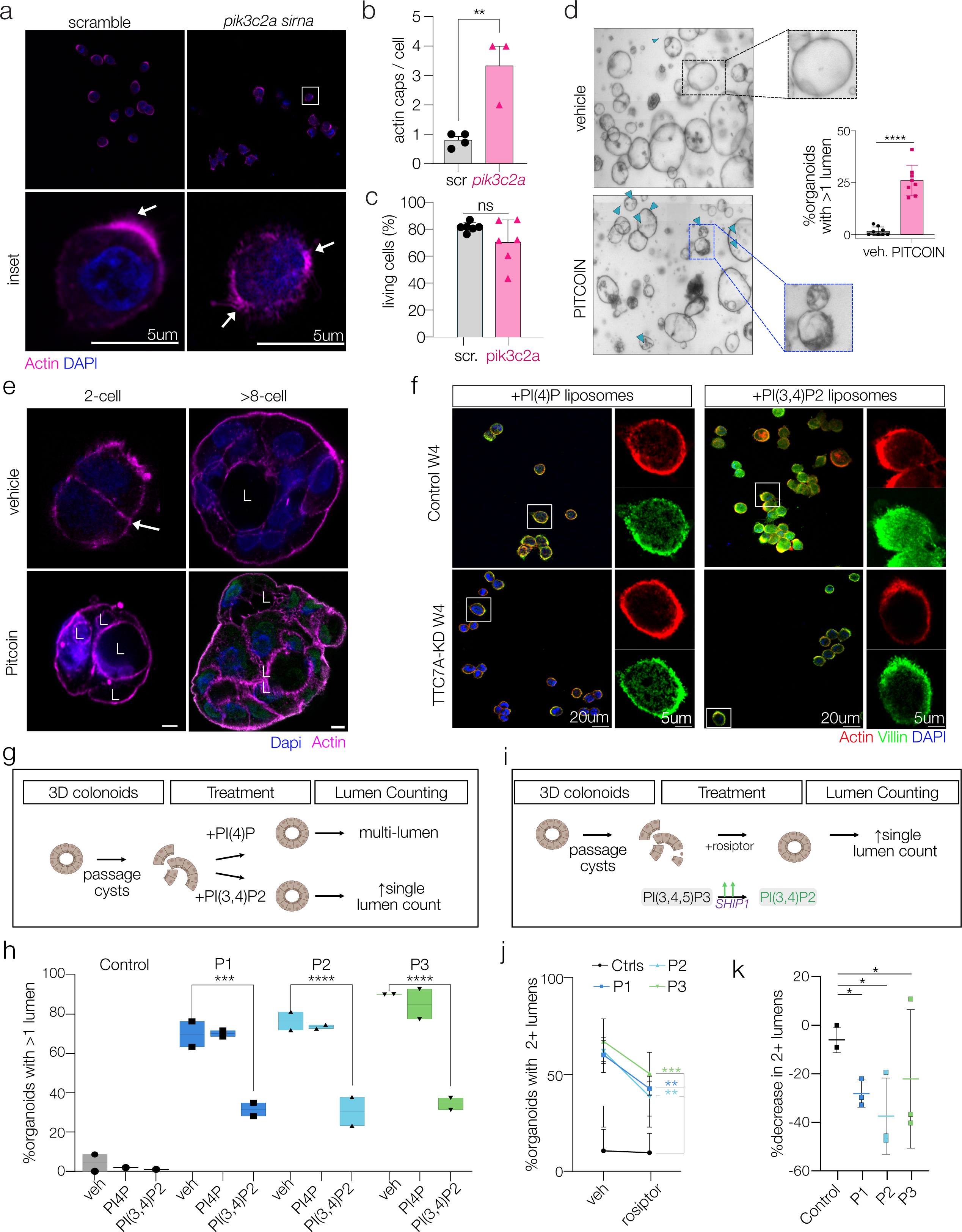
Abnormal lumen formation in TTC7A is recapitulated with loss of PIK3C2A and rescued with restoring PI(3,4)P2 levels. **a.** Overview images of WT-W4 cells treated with scrambled or *pik3c2a* siRNA and stained for Actin (magenta). Inset from overview. White arrows indicate actin cap formation in insets. **b.** Quantification of number of actin caps per cell with scrambled or *pik3c2a* siRNA represented as actin caps/cell. **c.** Viability of scrambled or *pik3c2a* siRNA W4 cells presented as bar graphs of FACS plots for calcein blue (+) cells. Represented as % living cells. **d.** Control colonoids treated with PITCOIN and quantification of organoids with >1 lumen. Inset indicate representative brightfield images with single, or >1 lumen per cyst. **e.** Immunofluorescent staining of control colonoids at the 2-cell or >8-cell stage, treated with vehicle or PITCOIN. Lumens indicated with L. **f.** Control-W4 and TTC7A-KD W4 cells treated with PI(4)P or PI(3,4)P2 liposomes stimulated with doxycycline. Inset shows representative images from each treatment. **g.** Schematic representation of colonoid liposome treatment strategy. **h.** Quantification of % of organoids with >1 lumen after treatment with liposomes+ vehicle, PI(4)P or PI(3,4)P2. **i.** Schematic representation of treatment with SHIP1 activator, rosiptor on colonoids. **j.** Quantification of changes in %of organoids with >1lumen post rosiptor treatment in control (average is n=3 images) or three patient colonoids. **k.** Representation of %decrease in secondary lumens post rosiptor treatment in control (average is n=3 patients) or three patient colonoids.

Since PIK3C2A function is likely required for single lumen formation, we assessed whether we could rescue the TTC7A patient phenotype in W4 cells by providing the enzymatic product of PIK3C2A, PI(3,4)P_2_. To do this we administered liposomes incorporating either PI(3,4)P_2_ or PI(4)P to both W4 and TTC7A-KD W4 cells. We found that addition of PI(3,4)P_2_ liposomes but not PI(4)P liposomes induced a significant increase in single-actin cap formation in TTC7A-KD W4 cells (Fig 6f). Next, we tested if addition of PI(3,4)P_2_ could rescue the abnormal lumen defect in patient derived colonoids (Fig 6g). We observed that treatment with PI(3,4)P_2_ liposomes could rescue the multiple-lumen phenotype in all three TTC7A^LOF^ colonoids, in contrast to PI(4)P which had no effect (Fig 6h, Supplementary 6f).

Since PI(3,4)P_2_ generation can be pharmacologically stimulated^37,38^, we investigated whether we could therapeutically rescue the multiple lumen phenotype seen in TTC7A-patients by activating SHIP1 (Fig 6i) with the small-molecule activator rosiptor. Treatment of TTC7A^LOF^ colonoids with rosiptor induced a significant decrease in the percentage of colonoids with secondary lumens (Fig 6j) compared to vehicle (Fig 6k), providing evidence that increasing PI(3,4)P_2_ is sufficient to rescue the epithelial phenotype seen in TTC7A patients.

## Discussion

An important morphological feature of many tissues is the presence of tubular structures comprising of a single layer of epithelial cells surrounding a central lumen. In the colon, formation of tubular crypt glands is critical for normal mucosal absorption, regeneration, and barrier function. Self-organization of these multi-cellular structures requires specification, localization and formation of polarized membrane domains to direct the site of the central lumen and organize key protein and lipid mediators. Here, we used TTC7A loss of function models to dissect the spatial and cell-state dependent temporal dynamics of epithelial lumen formation. Patients with loss of function mutations in the chaperone protein TTC7A exhibit a remarkable colonic crypt gland multi-lumen phenotype, which we show is consistently recapitulated in epithelial organoids, and suggestive of a key role for TTC7A in *de novo* lumen morphogenesis. We show that TTC7A is essential for directionally localizing and maintaining the apical membrane domain and juxtamembane PIPs, providing a framework for understanding how TTC7A-dependent cellular processes link to disease pathology. Our work identifies a novel chaperone function of TTC7A in lumen formation that is distinct from its previously established role as a plasma membrane protein and PI4KIIIΑ chaperone. Finally, we demonstrate that restoration of PI(3,4)P2 through exogenous delivery or pharmacological modulation rescues aberrant lumen formation.

Lumen formation has been investigated in a variety of in 3D epithelial culture systems and is dependent on the establishment of polarized membrane domains and organization of the apical membrane^2,8,9,39^. *In vivo* studies have shown that this reorganization of membrane proteins occurs via Rab4- or Rab11a-mediated vesicular trafficking to establish nascent apical membrane domains, which fuse to form a tubular structure spanning several cells^6,40,41^. Our data imaging the initial cell divisions of forming healthy colonoids demonstrate that epithelial cells develop multiple small apical membrane domains that then coalesce into a unified domain. This process mirrors developmental lumenogenesis previously observed in the zebrafish intestine as well in rat colonic crypt glands, where multiple independent lumens open and enlarge before resolving into a single continuous lumen.^6,42^ Lumen resolution appears to require the vesicular remodeling of contacts between adjacent lumens and subsequent lumen fusion in a distinct process separate from initial lumen opening.^6^ In TTC7A^LOF^ colonoids, however, multiple lumens persist past these initial cell divisions suggesting a failure of lumen coalescence, that is TTC7A dependent, and involves alteration of the vesicular remodeling process. Manipulation of TTC7A localization by directed chemically induced dimerization indicated that TTC7A function is required to direct apical membrane localization but only early during the process of cell polarization. Loss of TTC7A was associated with the persistence of multiple membrane sites associated with an apical identity suggesting a necessary role in directing apical domain location but not in membrane polarization *per se*.

Previous studies have outlined a role for TTC7A in a multi-protein complex with PI-kinases^21,25,43^. While TTC7A and its paralog TTC7B have been shown to associate with PI4KIIIα, our studies provide evidence for an additional role for TTC7A as a chaperone for the PI-kinase - PIK3C2A. The interaction between TTC7A and PIK3C2A was confirmed by co-immunoprecipitation experiments showing a strong association in control colonoids, which was significantly reduced in TTC7A^LOF^ cells indicating a role in the epithelial architectural manifestations seen in TTC7A patients. In addition to loss of association with PIK3C2A with TTC7A mutations, we did also observe reduced association with PI4KIIIΑ in mature, differentiated colonoids. We find that TTC7A-dependent association with PI4KIIIΑ does not have a major role in the process of apical membrane localization and glandular lumen formation but is critical for cell survival signaling^18^. Human loss-of-function mutations in PI4KIIIΑ are associated with a complex clinical phenotype that is most consistently characterized by neurological abnormalities but also variably by GI manifestations including inflammation and multiple intestinal atresia.^44–46^ The ability of TTC7A to chaperone both kinases, likely due to topological similarities in their alpha solenoid domains despite low sequence homology^26^, suggests that TTC7A has a dual role in epithelial biology, regulating survival pathways via interaction with PI4KIIIΑ and lumen formation through PIK3C2A.

A key component of the initial stages in lumen coalescence is remodeling by vesicular trafficking to and from developing apical membrane domains.^1,8^ Our analyses reveal that TTC7A and PIK3C2A localize to Rab11a-positive vesicles during a critical window when colonic epithelial cells specify and direct the nascent apical membrane. TTC7A is likely to be important in chaperoning PIK3C2A to the plasma membrane via directed trafficking. In this context, loss of TTC7A function leads to mis-directed trafficking of PIK3C2A in Rab11a-positive recycling endosomes resulting in persistence of multiple apical domains and loss of the remodeling that occurs early in lumen fusion. Perturbations in the Rab11 trafficking pathway have been shown to impair single lumen formation in other models, including zebrafish lumen formation, where Rab11- mediated trafficking is necessary for proper lumen coalescence.^6^ Consistent with a similar role in human colon crypt coalescence, Rab11^+^ vesicles without TTC7A, while capable of transporting cargo, fail to correctly direct cargo to a unified apical membrane, resulting in the delivery of apical components to multiple sites.

Interestingly, loss of TTC7A function reduces PI(3,4)P2 production without significantly affecting PI(4)P expression in colonoids, suggesting that compensatory mechanisms by other chaperones such as Fam126 and TMEM150 may maintain PI(4)P production in TTC7A patient cells.^22^ PI(3,4)P2 is generated by PIK3C2A therefore likely directs the identity of the nascent apical membrane domain with subsequent recruitment of effector proteins to coordinate membrane remodeling and junction formation during lumen initiation. Thus, direct modulation of PI(3,4)P2 rescues epithelial defect caused by TTC7A^LOF^ bypassing PIK3C2A.

There are currently very limited options for treatment of the intestinal epithelial defects seen in TTC7A patients.^18^ Mortality and morbidity remain extremely high and there remains a need for novel therapeutic approaches.^16^ We identified two therapeutic avenues that address the lumen morphogenic defects due to TTC7A dysfunction. First, we showed that exogenous administration of PI(3,4)P2, but not PI(4)P-liposomes, to TTC7A patient-derived colonoids effectively rescued the multiple-lumen phenotype, demonstrating a lipid-specific salvage effect. This strategy bypasses the requirement for functional TTC7A-PIK3C2A interactions by directly restoring the lipid at the site of action. The selective restoration of lumen coalescence by PI(3,4)P2 also further highlights the required specificity of phosphoinositide signaling in lumen formation. We also identified a small molecule approach using a pharmacological activator of SHIP1 that induces PI(3,4)P2 synthesis and restores apical membrane coalescence in patient-derived colonoids. This approach bypasses the need for TTC7A function by amplifying the residual activity of PIK3C2A, effectively compensating for the loss of its chaperone function. These studies therefore provide a a rationale for new therapeutic options targeted at TTC7A^LOF^-associated epithelial defects and further studies are needed to assess whether these may be viable for patients with TTC7A disease. In addition, given the critical role of phosphoinositide regulation by TTC7A and lipid kinases in tissue morphogenesis and maintenance of the epithelial barrier, developing targeted interventions that modulate their function could be a potential strategy for epithelial-directed therapies in IBD in general.

---

**Supplemental Figure 1:**
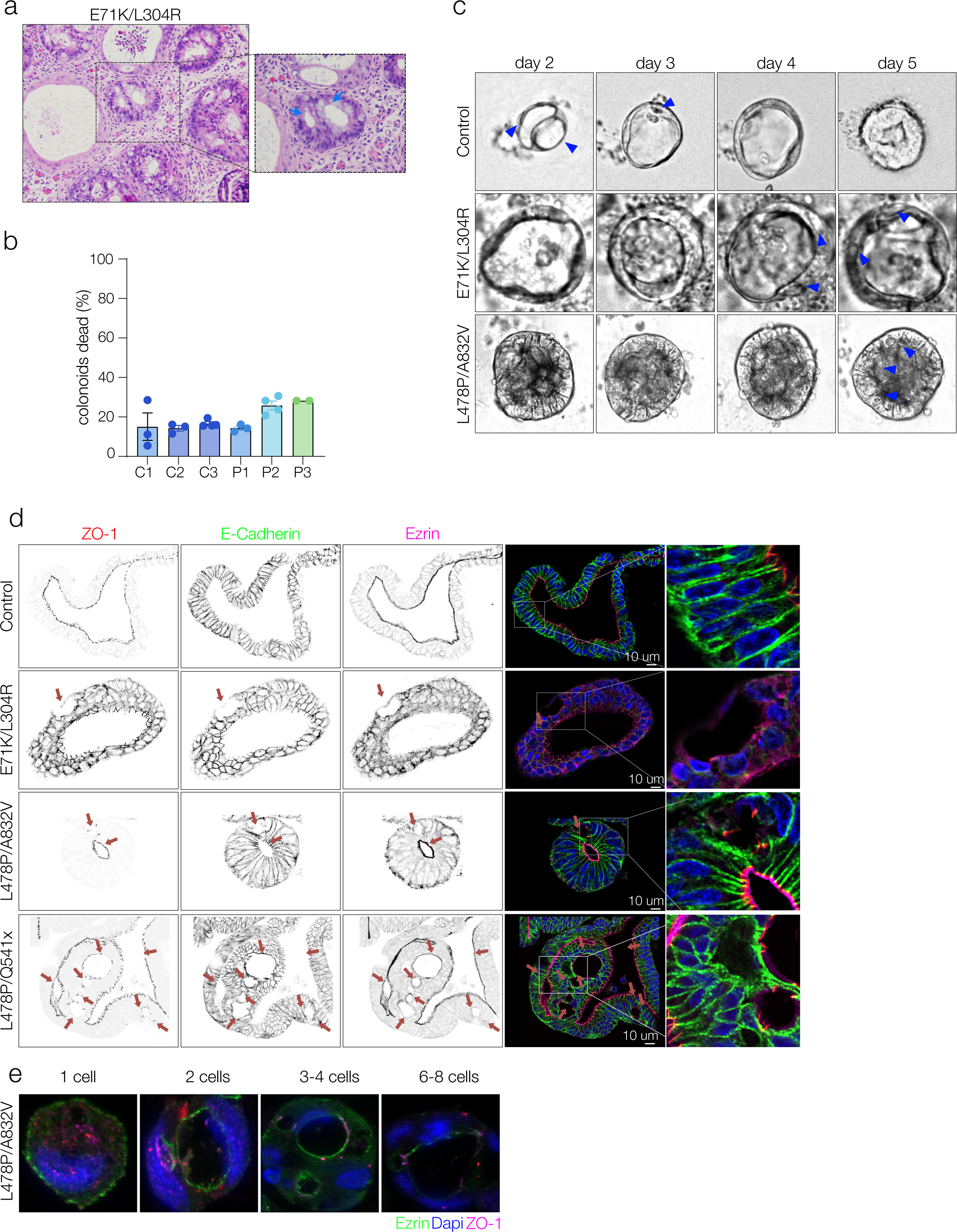
**a.** Histopathology of patient tissue showing crypt glands with multiple secondary lumens in a patient with TTC7A mutations (E71K/L304R). Inset with blue arrows highlight secondary lumen structures in crypt glands. **b.** Quantification of dead organoids in control and patient colonoids (n=3; data presented as one-way ANOVA corrected for multiple comparisons; no star = n.s., *<0.05, **<0.01, ***<0.001, ****<0.0001)). **c.** Brightfield time-lapse imaging of cyst development over 5 days, secondary lumens indicated by blue triangles. **d.** Representative confocal images with inverted micrographs on staining of ZO-1, Ezrin and E-Cadherin in control, and E71K/L304R, L478P/A832V, and L478P/Q541x colonoids. Insets show luminal expression of apical ezrin, junctional ZO-1 and basolateral E-Cadherin. **e.** Lumen development from single-cell to 8-cell stage in L478P/A832V patient derived colonoids. Ezrin in (green) and ZO-1 (red) marking the apical membrane.

**Supplemental Figure 2:**
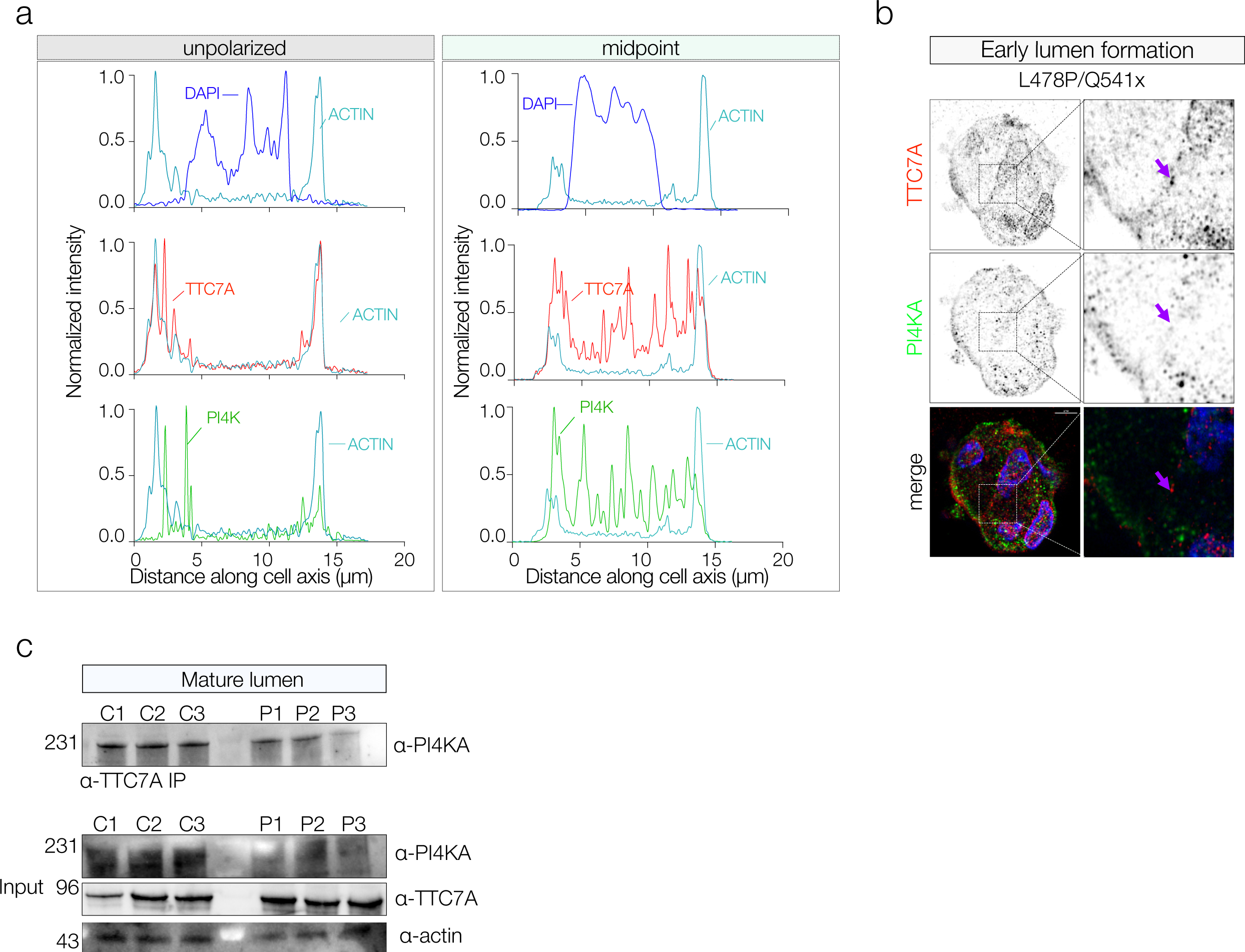
**a.** Representative plot profiles showing normalized intensity vs. cell length for the localization of Actin (teal) with DAPI (blue), TTC7A (red), and PI4KIIIΑ (green) in unpolarized, and polarized W4 cells. **b.** Representative confocal imaging of TTC7A^LOF^ colonoids showing endogenous TTC7A (red) and PI4KIIIΑ (green) subcellular localization in early lumens and in mature lumens. Dashed boxes indicate the zoomed insets of the lumen. Insets show diverging TTC7A and PI4KIIIΑ in early lumen formation. Grey-scale images are inverted black and white confocal images. **c.** Representative western blot on CO-IP TTC7A-pulldown probing for PI4KIIIΑ (top) in early lumen formation in three control and three patient colonoid lines. Data is representative of 6 experimental replicates. Western blot on whole cell lysates in three control and three patient colonoid lines probing for PI4KIIIΑ, TTC7A and actin. Data is representative of >6 experimental replicates.

**Supplemental Figure 3:**
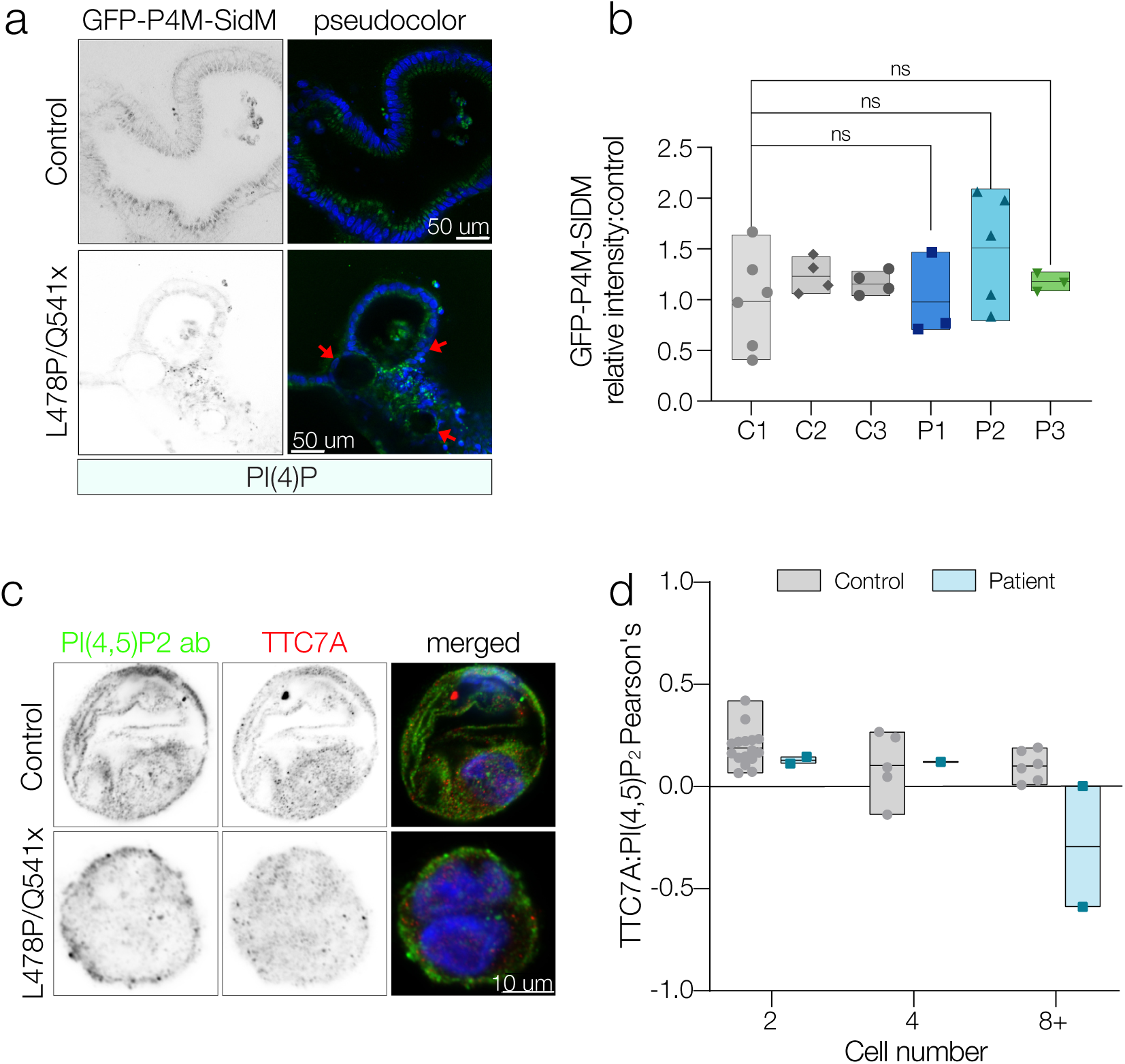
**a** Fluorescent imaging of GFP-P4M-SIDM probes in control and L478P/Q541x colonoids. **b.** Quantification of GFP-P4M-SIDM probes in control and patient colonoids. **c.** Immunostaining of PI(4,5)P2 (green) and TTC7A (red) in Control and L478P/Q541x colonoids. **d.** Pearsons correlation of TTC7A: PI(4,5)P2 staining. (n=3 controls, n=3 patients, ordinary two-way ANOVA with Fisher’s test)

**Supplemental Figure 4.**
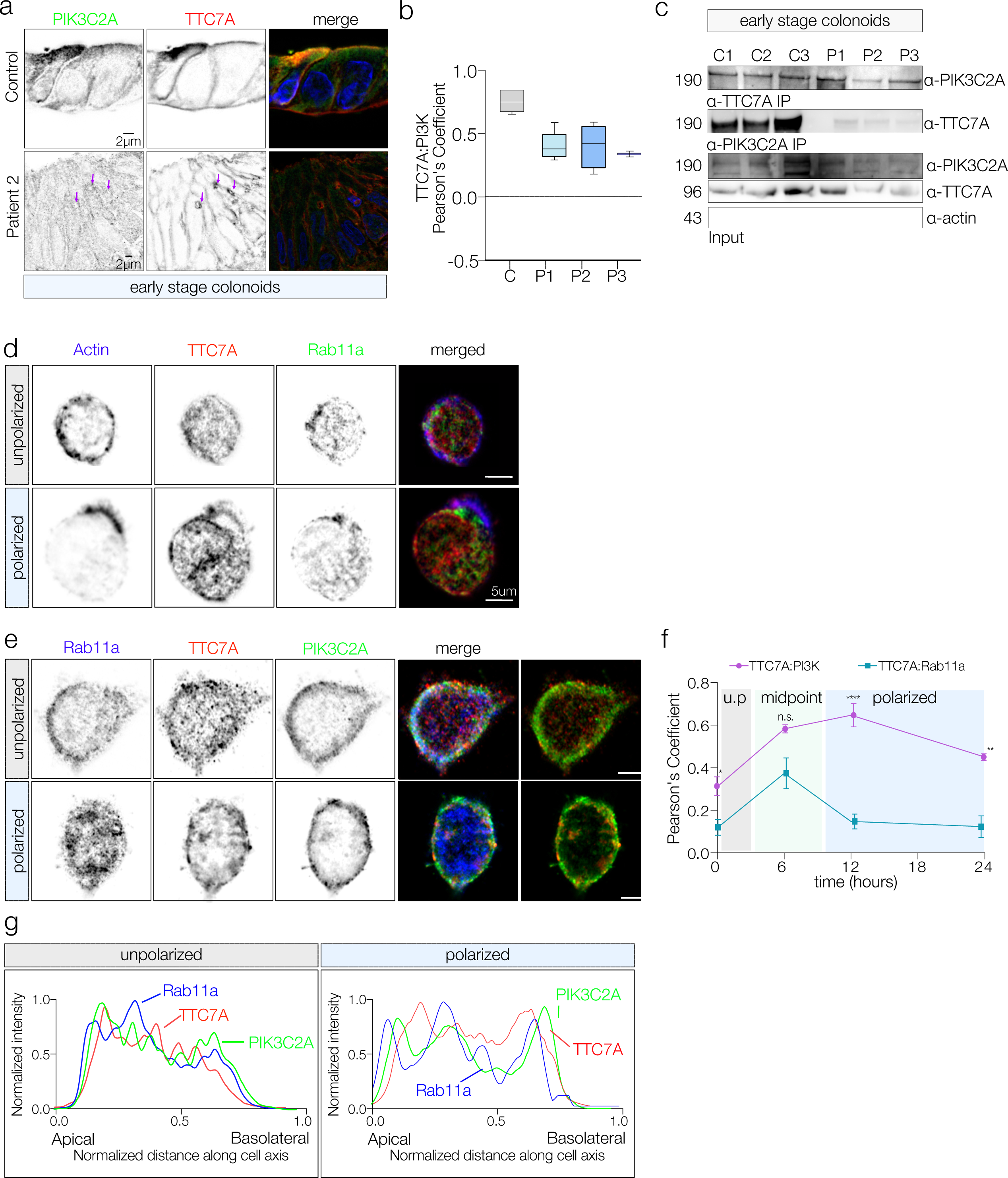
**a.** Representative confocal imaging of endogenous immunostaining of PIK3C2A (green) and TTC7A (red) in mature control vs. TTC7ALOF colonoids. Scale bars = 2uM. **b.** Pearson’s correlation for TTC7A and PIK3C2A localization in mature control (C) and TTC7ALOF colonoids (P1, P2, P3). (n=5 images per sample, n=3 patients, n=3 controls). **c.** CO-IP of TTC7A and PIK3C2A in mature colonoids and immunoblotting for TTC7A, PIK3C2A and Actin on the input samples, representative data from 5 experimental replicates. **d.** Immunostaining of Actin (blue) TTC7A (red) and Rab11a (green) at the unpolarized and polarized state of W4 cells. Scale bars = 5 uM. **e.** Immunostaining and confocal imaging on unpolarized and polarized W4 cells midpoint TTC7A (red), PIK3C2A (green), and Rab11a (blue), and merged TTC7A:PIK3C2A:Rab11a showing colocalization (white), TTC7A:PIK3C2A colocalization (yellow). Scale bars = 10um. **f.** Line plot for pearson’s Coefficient on WT-W4 for Rab11a:TTC7A (teal line) and TTC7A:PIK3C2A (magenta line) demonstrating average protein interactions across time, unpolarized (0-3hrs), midpoint (3-8hrs) and polarized (8-24hrs) cell states. **= p<0.001. One-way ANOVA with Dunnett’s multiple comparisons test. **g.** Representative plot profile of unpolarized and polarized W4 cells showing TTC7A (red), PIK3C2A (green), and Rab11a (blue).

**Supplemental Figure 5.**
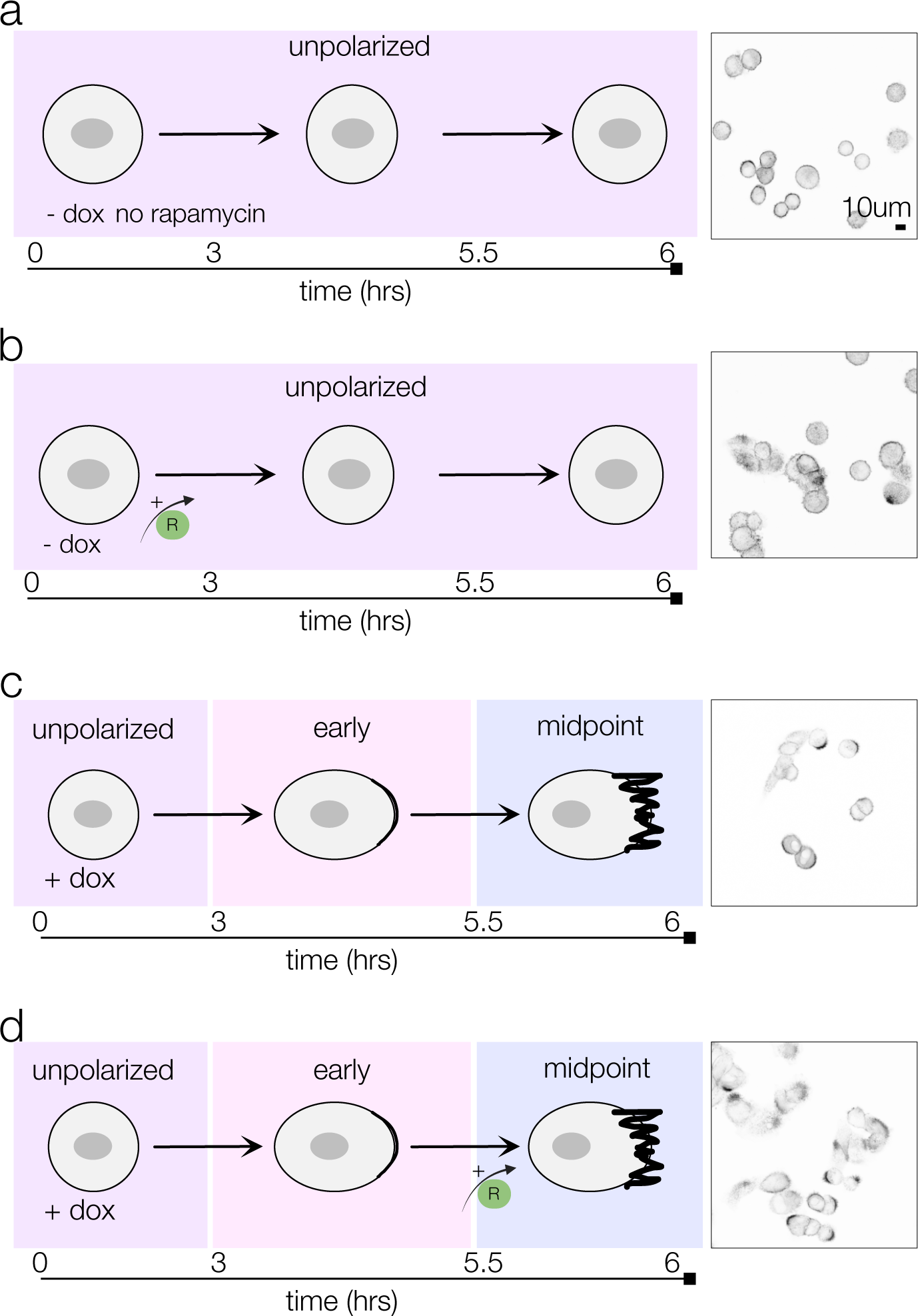
**a.** TTC7A-FKBP-PM-FRB W4 cells unpolarized with no rapamycin induced dimerization stained for actin. **b.** TTC7A-FKBP-PM-FRB W4 cells with rapamycin treatment for 6 hours stained for Actin **c.** TTC7A-FKBP-PM-FRB W4 cells with doxycycline for 6 hours but no rapamycin treatment stained for actin. **d.** TTC7A-FKBP-PM-FRB W4 cells with doxycycline treatment for 6 hours and rapamycin induced dimerization for 30 minutes stained for Actin.

**Supplemental Figure 6:**
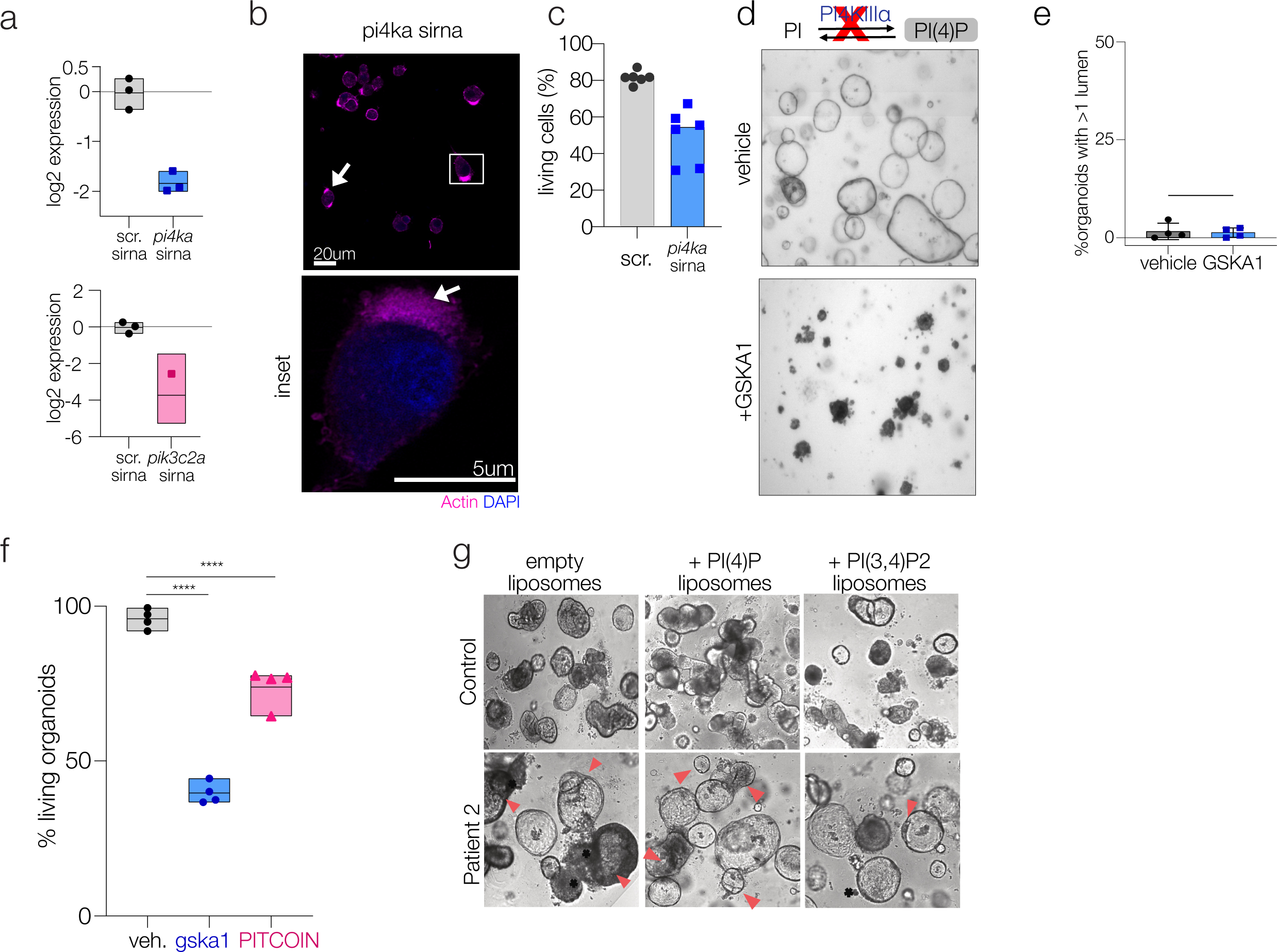
**a.** qPCR expression of PI4KIIIΑ or PIK3C2A after siRNA treatment. **b.** Representative actin staining in W4 cells treated with PI4KIIIΑ sirna. **c.** Viability of scrambled or *pi4ka* siRNA W4 cells presented as bar graphs of FACS plots for calcein blue (+) cells. Represented as % living cells **d.** Brightfield images of control colonoids treated with GSKA1. **e.** Quantification of GSKA1 treated colonoids with >1 lumen. **f.** Quantification of % living organoids post GSKA1 or PITCOIN treatment in control colonoids. **g.** Representative images of control and L478P/A832V patient colonoids treated with empty-liposomes, PI(4)P liposomes or PI(3,4)P2 liposomes.

## Methods

### Cell culture and drug treatments

LS174T:W4 cells were cultured according to the protocol developed by Baas et al^30^. Briefly, cells were routinely maintained in RPMI (Gibco 61870127), 100 μg/mL G-418 (Gibco 10131-035), 10 μg/mL blasticidin (Gibco A1113903), 20 μg/mL phleomycin (Invivogen, ANT-PH), and 10% tetracycline-free fetal bovine serum (FBS) (Takara Bio 631107). To induce brush border formation, doxycycline (10ug/ml) (Sigma 3072) was added. Caco2 and HEK293T cells were routinely maintained in DMEM (Fisher MT-10-013-CV) + 10% FBS + anti/anti. All cultures were maintained in humidified chamber, at 37°C with 5% CO_2_.

### Organoid culture

Colonic biopsy samples were obtained and cultured with methods modified from Sato et al.14 Briefly, crypts were dissociated from colonic biopsy samples obtained from a patient with TTC7A mutation or from a healthy control individual. Isolated crypts were suspended in Growth Factor Reduced Phenol Red Free Matrigel and plated as 30ul domes in a 24-well plate (Costar 3470). Media was replenished every other day.

Colonoid cultures were passaged with a modified mechanical passaging protocol as previously described^29^. Briefly, matrigel domes were washed with PBS, transferred to a conical tube, and pelleted in in matrigel. The PBS and free matrigel are aspirated and washed once more with PBS. A small portion of matrigel is left on top of the organoids, and organoids were passed through a P200 tip 50 times to fragment lines. Organoids were washed of dead cell debris, and replated in 30ul matrigel domes.

### Patient derived hematoxylin and eosin stain

H&E images were obtained from clinical colonic H&E biopsies.

### Lentivirus production

Lentiviral particles were prepared in HEK293 using standard protocols. HEK293T cells were transfected with the TTC7A-shRNA and psPAX2 and VSVG packaging plasmids with Lipofectamine 3000 (Thermo L3000008) in OptiMEM (Gibco 31-985-070). Transfection media was changed after 24 hours. Lentiviral particles were collected 72hours after transfection and concentrated using Lenti-X concentrator (Takara 631232) according to manufacturer’s protocol. Viral particles were quantified using Lenti-X GoStix Plus (Takara 631280) and volumetrically titrated to a transduction efficiency of >80% in HEK293T. Caco2 and LS174T:W4 cells were transduced with TTC7A-lenti particles using 10ug/ml Polybrene (Santa Cruz Biotechnology, sc-134220). A mixed population of cells was selected for using 10ug/ml puromycin (Sigma 540411- 25MG). TTC7A knockdown was confirmed by qPCR and western Blotting.

### RNA Isolation qPCR

Total RNA was extracted using Trizol/chloroform extraction with isopropanol and ethanol precipitations. cDNA was generated fromn 100ng RNA using iScript reverse transcription supermix (Bio-Rad). Primers were designed using Primer3 software^47^. qPCR was performed using SSO Advanced Universal SYBR Green Supermix (Bio-Rad) on an Applied Biosystems Real-Time PCR instrument. Relative gene expression was calculated using the ΔΔCt method and graphed as Log2 expression.

### Immunocytochemistry

#### Cell lines and organoid single cells

For whole mount organoid stains, organoids were transferred to a 15ml conical in matrigel, washed once with ice cold 2mM EDTA, and centrifuged at 300xg for 5 minutes. EDTA was aspirated and replaced with fresh ice-cold Cell recovery solution (Corning 354253) for 30 minutes at 4°C on a rotator until Matrigel was fully dissolved. Organoids were pelleted and fixed with 4%PFA and 1% Glutaraldehyde for 2hrs at RT with constant agitation. Organoids were allowed to settle by gravity, fixative was discarded, and pellet was washed with PBS-1%BSA (Sigma A6003-25G)-0.1%-mouse IgG (Invitrogen™ R37621), followed by a 10- minute wash in 0.2M Glycine-PBS. Glycine was removed and cells were permeabilized with PBS- 1% TritonX100 overnight at 4°C with constant agitation. Cells were then antigen retrieved on a heat plate set to 95°C for 10 minutes in Tris HCl pH 9.0 and allowed to cool for 10 additional minutes. After retrieval, cells were washed thrice with PBS-0.05%Tween-1%BSA-0.1%mouse IgG and blocked with 10%FBS-1%mouse IgG-in PBS-1%Tween. Primary and secondary antibodies were stained overnight in PBS-0.1%BSA-0.05%Tween. After removal of secondary antibody solution, cells were incubated with DAPI (1:10,000 in dH2O) (Thermo 62248) for 20 minutes. Organoids were washed in PBS-0.1%BSA-0.05%Tween 5 times and cleared through sequential H2O, H2O: Glycerol, 100% Glycerol washes. Organoids were mounted on depressed slides with Prolong Diamond Antifade (ThermoFisher P36970).

### Paraffin embedded

Organoids were isolated from matrigel and resuspended in 4% Agarose/30% Sucrose. Agarose blocks were fixed in paraformaldehyde and processed by the BIDMC histology core. 5um sections were deparaffinized and antigen retrieved in Tris-HCl, pH 9.0, and stained as described above.

### Antibodies for immunostaining and western blotting

| Target / Probe | Supplier | Catalog # | Application | Dilution factor | Type of probe |
| --- | --- | --- | --- | --- | --- |
| Acti-Stain 555 Phalloidin | Cytoskeleton | (Cat. # PHDH1-A) | IF | 1:500 | Fluorescent probe |
| Ezrin-Alexa fluor 488 | Abcam | AB198520 | IF | 1:250 | Direct conjugate |
| E-Cadherin – 647 | BD Biosciences | 560064 | IF | 1:250 | Direct conjugate |
| ZO-1 – 555 | Thermofisher | MA3-39100-A555 | IF | 1:100 | Direct conjugate |
| TTC7A | LS Bio | LS-C799928 | IF / CO-IP / WB | 1:100 / 1ug / 1:1000 | Primary Ab |
| PI4KIIIA | Proteintech | 12411-1-AP | IF / CO-IP / WB | 1:100 / 1ug / 1:500 | Primary Ab |
| PIK3C2A | Novus | NB300-855 | IF / WB | 1:100 / 1:500 | Primary AB |
| PIK3C2A | LS Bio | LS-C668683-100 | CO-IP | 1ug | Primary AB |
| Anti-PI(4,5) P2 IgM | Echelon Biosciences | Z-PO45 | IF | 1:100 | Primary AB |
| Anti-PI(3,4)P2 IGG | Echelon Biosciences | Z-P034B | IF | 1:100 | Primary AB |
| GAPDH | Thermofisher | MA515738HRP | WB | 1:1000 | Direct Conjugate |
| TidyBlot Western Blot Detection Reagent:HRP | Bio Rad | **STAR209PA** | CO-IP | 1:5000 | Secondary AB |
| Anti-mouse IgG, HRP-linked Antibody #7076 | Cell Signaling | **7076S** | WB | 1:5000 | Secondary AB |
| Anti-rabbit IgG, HRP-linked Antibody #7074 | Cell Signaling | **7074S** | WB | 1:5000 | Secondary AB |
| Donkey Anti Goat HRP | Fisher | **705-035-003** | WB | 1:5000 | Secondary AB |
| Alexa Fluor® 647 AffiniPure Donkey Anti-Mouse IgG (H+L) | Jackson ImmunoResearch | **715-605-151** | IF | 1:500 | Secondary AB |
| Alexa Fluor® 555 AffiniPure™ Donkey Anti-Mouse IgG (H+L) | Jackson ImmunoResearch | **715-565-150** | IF | 1:500 | Secondary AB |
| AffiniPure™ Donkey Anti-Rabbit IgG (H+L) | Jackson ImmunoResearch | **711-545-152** | IF | 1:500 | Secondary AB |
| AffiniPure™ Donkey Anti-Rabbit IgG (H+L) | Jackson ImmunoResearch | **711-475-152** | IF | 1:500 | Secondary AB |
| AffiniPure™ Donkey Anti-Goat IgG (H+L) | Jackson ImmunoResearch | **711-565-147** | IF | 1:500 | Secondary AB |
| Rab11a | Goldenring Lab | **VU-67** | IF | 1:500 | Primary AB |
| TTC7B | Abcam | ab193361 | WB | 1:1000 | Primary AB |
| Villin | SCBT | sc-58897 AF546 | IF | 1:50 | Direct Conjugate |
| Mem-glow | Cytoskeleton Inc. | MG07 | IF | 1:100 | Direct conjugate |

### Image acquisition and analysis

Confocal images were captured on both a Zeiss 880 + Airyscan and an Abberior StedyCon system (Abberior Instruments GmbH, Göttingen, Germany). Post-image processing for images captured on the StedyCon was done using Huygens Pro (Scientific Volume Imaging, Hilversum, Netherlands). For 3D cysts, all images were obtained at the medial plane of the cyst when possible. All analysis was carried out in FIJI using built-in ROI, measurement, and colocalization tools (^48^). Plot profiles were generated by averaging three rectangle sections across the cell.

Data were normalized to the maximum intensity per channel. Brightfield images of organoids were captured in 24-well plates (Corning Costar). Images were acquired using a Cytation 5 reader with Gen5 software. Custom imaging protocols were developed to capture the entire well contents across three z-planes. The Gen5 software’s built-in settings were used for image stitching and focus stacking. Quantitative analysis was performed using the cell counter plug-in in FIJI (ImageJ).

### Human Protein Atlas

Protein expression data were analyzed using R (version not specified) with tidyverse and ggplot2 packages. Raw immunohistochemistry (IHC) data were obtained from the Human Protein Atlas database and imported as a tab-separated values file. Analysis focused on the expression patterns of TTC7A and TTC7B genes across four gastrointestinal tissues: duodenum, small intestine, colon, and rectum. Expression levels were visualized using a ggplot2 heatmap with high, low and undetected values originating from HPA IHC data.

### Western Blot and Co-Immunoprecipitation

For Co-Immunoprecipatation, cells were solubilized by resuspension in lysis buffer^24^ (150mM NaCl, 50mM HEPES, 1% Triton-X100, 10% Glycerol, 1mM MgCl2, 1.0mM EGTA, Pierce Protease phosphatase inhibitor without EDTA (A32965)), passed through a 28G syringe four times, and incubated on ice for 10 min. Insoluble debris was pelleted at 12,000 x g for 30 min. The supernatant was harvested for input blots. For CO-IPs, 75ul of lysate was incubated with 1ug of antibody for three hours at RT, then 100ul of prewashed 50% bead mixture (Sigma Aldrich GE17- 0618-01) was added and incubated overnight at 4°C with constant rotation. Beads were allowed to settle by gravity, the supernatant discarded, and beads were washed 3x10minutes with lysis buffer. NuPAGE LDS (NP0008) + 1% DTT (Sigma D0632-1G) was added to each sample and boiled at 95°C for 5 minutes, immediately put on ice, and spun at 1200 RCF for 2 minutes. 20ul of lysate was added to a 4-12% Bis Tris Gel (NP0321BOX) and run at 80mAmp for 1 hour with NuPage MOPS SDS running buffer (NP0001).

Western blots were performed using standard protocols. Briefly, organoids or cells were removed completely from matrigel, and resuspended in lysis buffer (150mM NaCl, 50mM HEPES, 1% Triton-X100, 10% Glycerol, 1mM MgCl2, 1.0mM EGTA, Pierce protease phosphatase inhibitor without EDTA) or RIPA buffer (Boston Bio Products BP-115-500). Cells were passed through a 28G needle and incubated on ice for 10 minutes. Insoluble debris was pelleted at 12,000 x g for 30 minutes, supernatants collected, and mixed with NuPage LDS + DTT to a concentration of 10ug/ul. 20ul of lysate was added to a 4-12% BisTris gel, and run for 1 hour at 80mAmp in NuPage MOPS SDS buffer + 1x Antioxidant (NP0005).

All blots were transferred using an iBlot3 to PVDF membranes (Invitrogen IB34001). Blots were blocked with 5% skim milk in TBS-0.5%tween for 1 hour at room temperature. Primary antibodies were added to the blot at a 1ug/ml concentration in TBS-0.05% tween, and incubated overnight at 4°C. Blots were washed with TBS-0.5%Tween for 3x5 minute washes, and incubated with secondary antibodies at a 1:10,000 concentration for 1hr at RT. Blots were washed for 3x5minutes and imaged on a ChemiDoc with Pierce Femto (Fisher 34096).

For input, blots were briefly stripped with 0.2N NaoH for 2minutes at RT, washed, reblocked and probed with GAPDH-HRP (Fisher MA515738HRP) for 1hr at RT.

### Phosphoinositide probe transfection

The following probes were used for these studies: GFP-C1-PLCdelta-PH was a gift from Tobias Meyer (Addgene plasmid # 21179); NES-EGFP-cPHx3 was a gift from Gerry Hammond (Addgene plasmid # 116855); Grp1-PH pEGFP-C1 was a gift from Mark Lemmon (Addgene plasmid # 71378); GFP-P4M-SidM was a gift from Tamas Balla (Addgene plasmid # 51469), and 3xFlag- PI4KIIIα was a gift from Pietro De Camilli (PMC5748228).

Cells were transfected using Lipofectamine 3000 according to the manufacturer’s protocol. Organoids were transfected with liposomes composed of Lipofectamine RNAiMax (13778030) and P3000. Briefly, 1ug of dna is incubated with P3000 reagent in Optimem, while RNAiMAX is equilibrated in optimem. Equal volumes of P3000-DNA and RNAiMAX are mixed and incubated for 15 minutes at RT with rotation. Organoid pellets are resuspended in DNA-liposome mixture and spun at 300RCF for 30 minutes at 37C. After 30 minutes, all but 75ul of DNA-liposome mixture is removed from the organoid pellets, organoids are resuspended in Matrigel, and plated in 24- well glass bottom plates. Matrigel is allowed to polymerize at 37C for 15 minutes, and organoids are fed with plating media. After 24 hours, plating media is replaced with expansion media. Fluorescence is observable between 48hours and 72 hours post transfection, and cells are viable for up to 5 days post transfection.

### Live Cell imaging

Live cell imaging was performed using a Zeiss LSM880 with Airyscan detection. For adherent cell imaging, cells expressing GFP-tagged proteins were cultured on glass-bottom plates (CellVis). Prior to imaging, cells were washed twice with pre-warmed PBS, followed by the addition of live-cell imaging buffer.

For organoid imaging, samples were embedded in 50% Matrigel and plated on glass-bottom plates (CellVis). At the time of imaging, organoid domes were washed three times with PBS and stained with either Hoechst (1:10,000 dilution) or MemGlow for 15 minutes at 37°C in a 5% CO₂ environment. Following incubation, domes were washed three times with PBS, and live-cell imaging buffer was added. All imaging of PI-probes was performed under live-cell conditions.

### FK506 binding protein (FKBP) - FKBP–rapamycin binding (FRB) experiments

TTC7A-FKBP construct was designed with SeqBuilderPro (DNASTAR) and purchased from Twist-Bioscience. Full length sequence is available upon request. Plasma membrane PM-FRB-CFP was a gift from Tamas Balla (Addgene plasmid # 67517). mCherry-FKBP-PI4KIIIΑC1001 was a gift from Gerry Hammond (Addgene plasmid # 139311).

LS174T-W4 cells were seeded on a 24-well glass bottom plate (Cellvis: P24-1.5H-N) with a density of 0.1 x 10^6^ per well. On the following day, cells were transfected with TTC7A-FKBP using lipofectamine 3000 reagent (Invitrogen: L3000015) according to the manufacturer’s protocols. ∼12 hours post-transfection, cells were subjected to the following conditions before fixing using 4% Paraformaldehyde and 1% Glutaraldehyde in PBS; (1) no rapamycin/ no doxycycline (2) no rapamycin/ 6 h doxycycline (3) no rapamycin/ 24 h doxycycline (4) 5 min rapamycin/ no doxycycline (5) 5 min rapamycin/ 6 h doxycycline (6) 5 min rapamycin/ 24 h doxycycline (7) 3 h rapamycin/ no doxycycline (8) 3 h rapamycin/ 6 h doxycycline (9) 3 h rapamycin/ 24 h doxycycline. Final concentrations of rapamycin and doxycycline were 5 nM and 10 µg/ml, respectively. Control experiments were conducted in cells without TTC7A-FKBP expression in similar conditions as before.

### Liposome preparation and composition

Liposomes were prepared with modifications to previously described methods^49^ (elife 91345). Acceptor liposomes were formed consisting of 2%NBD-PS 10% DOPE, 86%DOPC. Donor liposomes were composed of DOPC + 5% PI(4)P, PI(3,4)P2 or DOPC alone. Lipids were mixed in a glass container, and solvent was evaporated using a speedvac to create a film. Lipids were then freeze-dried overnight under vacuum. Next, lipid films were rehydrated in 50mM HEPES, 120mM potassium acetate and 1mM magnesium chloride, with rotation, for 30 minutes. Rehydrated lipid solutions were then freeze thawed on dry ice for 5 cycles of 5- minutes at each temperature. Lipids were extruded using an Avanti MiniExtruder with a 100nm filter prior to assays.

#### Inhibitor studies

Organoids were treated with the following chemicals: GSKA1 (Cayman 34502), SHIP1 (EMD-565835), and PITCOIN3 (a gift from Volker Haucke PMID:36109648).

#### Flow cytometry

LS174T:W4 cells were trypsinized, and resuspended in FACS buffer (1× PBS supplemented with 1% BSA and 2mM EDTA, with Y27632). For viability assessment, cells were stained with a dual-labeling approach using Calcein Red-Orange AM (50µM working solution, Invitrogen C34851) and Zombie Violet (Biolegend 423113) fixable dye. Staining was performed by incubating cells with Calcein (2µl/ml of 50µM solution) and Zombie dye (1:500 dilution) in FACS buffer for 15-30 minutes at 4°C, protected from light. Following staining, cells were washed with Biolegend Cell Staining Solution (Cat. 420201), and filtered through a 40µm filter cap FACS tube immediately prior to analysis. Analysis was performed on a BD Fortessa flow cytometer and data were analyzed using FlowJo.

#### siRNA studies

Predesigned Silencer Select siRNA constructs were used for interference studies. Constructs for PI4KIIIΑ (ThermoFisher, AM51331), PIK3C2A (ThermoFisher, 4390824, Assay ID 10508), and Efr3a (ThermoFisher, AM16074, ID 219474) were purchased from ThermoFisher. siRNA constructs were transiently transfected into LS174T:W4 cells using Lipofectamine RNAiMAX Transfection Reagent (ThermoFisher 13778030) according to the manufacturer’s protocol, 24 hours prior to the induction of polarization.

#### Protein modeling

The partial crystal structure of PI4KIIIΑ:TTC7B (PDB: 6BQ1) (PMID: 29229838) was downloaded the RCSB Protein Data Bank and visualized using Protean3D version 17 (DNAStar). Full-length predicted structures for TTC7A (PDB: AF-Q9ULT0-F1), PI4KIIIΑ (PDB: AF-P42356-F1), Fam126 (PDB: AF-Q9BYI3-F), and PIK3C2A (PDB: AF-O00443-F1) were downloaded from AlphaFold. Alignment of full-length AlphaFold structures against the partial crystal structures was performed using jFatCat rigid-body alignment for visualization of the full-length complex. Patient mutations in TTC7A were visualized using Protean3D version 17 (DNAStar).

#### Study approval

The Institutional Review Boards of Boston Children’s Hospital, Seattle Children’s Hospital, and Texas Children’s Hospital approved this study (BCH IRB P00027983, SCH IRB STUDY00002767, and TCH IRB H-8226) and informed consent/assent was given in accordance with the Declaration of Helsinki.

#### Statistics and data presentation

Statistical significance was calculated by GraphPad Prism software. Custom schema were designed using Affinity Designer software.

## Data availability

All data supporting the findings of this study are available within the paper and its supplementary information or are available upon request from the corresponding author.

## Acknowledgements

We thank the patients and their families who participated in this study, as well as members of the Boston Children’s Congenital Enteropathy Program.

## Author information

Dhanushan Wijayaratna and Geordie Emberling contributed equally.

## Contributions

Conceptualization: KBG JRT

Methodology: KBG, JL, JRT

Validation: DW, GE, LG

Investigation – KBG, DW, GE, LG

Provided samples and clinical information – HBZ, DW, LBK

Writing - original draft - KBG

Writing – review and editing – AM, SBS, KR, KBG, JRT

Project administration - JRT

Funding acquisition: KBG, WIL, JRT

Supervision: SBS, KR, WIL & JRT

## Ethics declarations

Competing interests: SBS declares the following interests: Scientific advisory board participation for Pfizer, Merck, Sonoma Biotherapeutics, Spyre Therapeutics, Biolojic Design, Trex Bio.

## REFERENCES

1. Bugda Gwilt, K. & Thiagarajah, J. R. Membrane Lipids in Epithelial Polarity: Sorting out the PIPs. Front. Cell Dev. Biol. 10, 893960 (2022).

2. Martin-Belmonte, F. et al. PTEN-mediated apical segregation of phosphoinositides controls epithelial morphogenesis through Cdc42. Cell 128, 383–397 (2007).

3. Mathan, M., Moxey, P. C. & Trier, J. S. Morphogenesis of fetal rat duodenal villi. Am. J. Anat. 146, 73–92 (1976).

4. Gassama-Diagne, A. et al. Phosphatidylinositol-3,4,5-trisphosphate regulates the formation of the basolateral plasma membrane in epithelial cells. *Nat*. Cell Biol. **8**, 963–970 (2006).

5. Ng, A. N. Y. et al. Formation of the digestive system in zebrafish: III. Intestinal epithelium morphogenesis. Dev. Biol. 286, 114–135 (2005).

6. Alvers, A. L., Ryan, S., Scherz, P. J., Huisken, J. & Bagnat, M. Single continuous lumen formation in the zebrafish gut is mediated by smoothened-dependent tissue remodeling. Dev. Camb. Engl. 141, 1110–1119 (2014).

7. Bryant, D. M. et al. A Molecular Switch for the Orientation of Epithelial Cell Polarization. Dev. Cell 31, 171–187 (2014).

8. Román-Fernández, Á. et al. The phospholipid PI(3,4)P2 is an apical identity determinant. Nat. Commun. 9, (2018).

9. Jewett, C. E. & Prekeris, R. Insane in the apical membrane: Trafficking events mediating apicobasal epithelial polarity during tube morphogenesis. Traffic Cph. Den. (2018) doi:10.1111/tra.12579.

10. Zegers, M. M. P., O’Brien, L. E., Yu, W., Datta, A. & Mostov, K. E. Epithelial polarity and tubulogenesis in vitro. Trends Cell Biol. 13, 169–176 (2003).

11. Rodriguez-Boulan, E. & Macara, I. G. Organization and execution of the epithelial polarity programme. Nat. Rev. Mol. Cell Biol. 15, 225–242 (2014).

12. Fölsch, H. The building blocks for basolateral vesicles in polarized epithelial cells. Trends Cell Biol. 15, 222–228 (2005).

13. Takahashi, S. et al. Rab11 regulates exocytosis of recycling vesicles at the plasma membrane. J. Cell Sci. 125, 4049–4057 (2012).

14. Mayinger, P. Phosphoinositides and vesicular membrane traffic. Biochim. Biophys. Acta 1821, 1104–1113 (2012).

15. Buckley, S. et al. Patients with TTC7A Deficiency Causing Multiple Intestinal Atresia with Combined Immunodeficiency (MIA-CID) Can Survive Well Beyond Infancy: a multicentre case series. Transplantation 101, S34 (2017).

16. Jardine, S., Dhingani, N. & Muise, A. M. TTC7A: Steward of Intestinal Health. Cell. Mol. Gastroenterol. Hepatol. 7, 555–570 (2018).

17. Dannheim, K. et al. Pediatric Gastrointestinal Histopathology in Patients With Tetratricopeptide Repeat Domain 7A (TTC7A) Germline Mutations: A Rare Condition Leading to Multiple Intestinal Atresias, Severe Combined Immunodeficiency, and Congenital Enteropathy. Am. J. Surg. Pathol. 46, 846–853 (2022).

18. S, J., et al. Drug Screen Identifies Leflunomide for Treatment of Inflammatory Bowel Disease Caused by TTC7A Deficiency. Gastroenterology 158, (2020).

19. Audhya, A., Foti, M. & Emr, S. D. Distinct roles for the yeast phosphatidylinositol 4-kinases, Stt4p and Pik1p, in secretion, cell growth, and organelle membrane dynamics. Mol. Biol. Cell 11, 2673–2689 (2000).

20. Nakatsu, F. et al. PtdIns4P synthesis by PI4KIIIα at the plasma membrane and its impact on plasma membrane identity. J. Cell Biol. 199, 1003–1016 (2012).

21. D, B., C, S., A, A., S, W. & Sd, E. Assembly of the PtdIns 4-kinase Stt4 complex at the plasma membrane requires Ypp1 and Efr3. J. Cell Biol. 183, (2008).

22. Chung, J., Nakatsu, F., Baskin, J. M. & De Camilli, P. Plasticity of PI4KIIIα interactions at the plasma membrane. EMBO Rep. 16, 312–320 (2015).

23. Baskin, J. M. et al. The leukodystrophy protein FAM126A (hyccin) regulates PtdIns(4)P synthesis at the plasma membrane. Nat. Cell Biol. 18, 132–138 (2016).

24. Lees, J. A. et al. Architecture of the human PI4KIIIα lipid kinase complex. Proc. Natl. Acad. Sci. U. S. A. 114, 13720–13725 (2017).

25. Avitzur, Y. et al. Mutations in tetratricopeptide repeat domain 7A result in a severe form of very early onset inflammatory bowel disease. Gastroenterology 146, 1028–1039 (2014).

26. Lees, J. A. et al. Architecture of the human PI4KIIIα lipid kinase complex. Proc. Natl. Acad. Sci. U. S. A. 114, 13720–13725 (2017).

27. Culbreath, K. et al. Intestinal atresias and intestinal failure in patients with TTC7A mutations. J. Pediatr. Surg. Case Rep. 80, 102247 (2022).

28. Tissue-based map of the human proteome | Science. https://www.science.org/doi/10.1126/science.1260419.

29. Gwilt, K. B. & Thiagarajah, J. R. Overcoming problematic growth phenotypes in organoids from patients with monogenic GI disease. (2023).

30. Baas, A. F. et al. Complete Polarization of Single Intestinal Epithelial Cells upon Activation of LKB1 by STRAD. Cell 116, 457–466 (2004).

31. Del Campo, C. M. et al. Structural basis for PI(4)P-specific membrane recruitment of the Legionella pneumophila effector DrrA/SidM. Struct. Lond. Engl. 1993 22, 397–408 (2014).

32. Stauffer, T. P., Ahn, S. & Meyer, T. Receptor-induced transient reduction in plasma membrane PtdIns(4,5)P2 concentration monitored in living cells. Curr. Biol. CB 8, 343–346 (1998).

33. Goulden, B. D. et al. A high-avidity biosensor reveals plasma membrane PI(3,4)P2 is predominantly a class I PI3K signaling product. J. Cell Biol. 218, 1066–1079 (2019).

34. P, V., B, T., T, R. & T, B. Rapidly inducible changes in phosphatidylinositol 4,5-bisphosphate levels influence multiple regulatory functions of the lipid in intact living cells. J. Cell Biol. 175, (2006).

35. Liu, L., Chen, L., Chung, J. & Huang, S. Rapamycin inhibits F-actin reorganization and phosphorylation of focal adhesion proteins. Oncogene 27, 4998–5010 (2008).

36. Lo, W.-T. et al. Development of selective inhibitors of phosphatidylinositol 3-kinase C2α. Nat. Chem. Biol. 19, 18–27 (2023).

37. Feng, Y. et al. Effective inhibition of miR-330/SHIP1/NF-κB signaling pathway via miR-330 sponge repolarizes microglia differentiation. Cell Biol. Int. 45, 785–794 (2021).

38. PubChem. Rosiptor. https://pubchem.ncbi.nlm.nih.gov/compound/76965484.

39. Martín-Belmonte, F. et al. Cell Polarity Dynamics Controls the Mechanism of Lumen Formation in Epithelial Morphogenesis. Curr. Biol. CB 18, 507–513 (2008).

40. Madara, J. L., Neutra, M. R. & Trier, J. S. Junctional complexes in fetal rat small intestine during morphogenesis. Dev. Biol. 86, 170–178 (1981).

41. Bagnat, M., Cheung, I. D., Mostov, K. E. & Stainier, D. Y. R. Genetic control of single lumen formation in the zebrafish gut. Nat. Cell Biol. 9, 954–960 (2007).

42. Madara, J. L., Neutra, M. R. & Trier, J. S. Junctional complexes in fetal rat small intestine during morphogenesis. Dev. Biol. 86, 170–178 (1981).

43. Bigorgne, A. E. et al. TTC7A mutations disrupt intestinal epithelial apicobasal polarity. J. Clin. Invest. 124, 328–337 (2014).

44. Saettini, F. et al. Biallelic PI4KA Mutations Disrupt B-Cell Metabolism and Cause B-Cell Lymphopenia and Hypogammaglobulinemia. J. Clin. Immunol. 45, 15 (2024).

45. Verdura, E. et al. Biallelic PI4KA variants cause a novel neurodevelopmental syndrome with hypomyelinating leukodystrophy. Brain 144, 2659–2669 (2021).

46. Salter, C. G. et al. Biallelic PI4KA variants cause neurological, intestinal and immunological disease. Brain J. Neurol. 144, 3597–3610 (2021).

47. Kõressaar, T. et al. Primer3_masker: integrating masking of template sequence with primer design software. Bioinforma. Oxf. Engl. 34, 1937–1938 (2018).

48. Schindelin, J., et al. Fiji: an open-source platform for biological-image analysis. Nat. Methods 9, 676–682 (2012).

49. Cabukusta, B. et al. The ORP9-ORP11 dimer promotes sphingomyelin synthesis. eLife 12, RP91345 (2024).

